# Psychological stress disrupts intestinal epithelial cell function and mucosal integrity through microbe and host-directed processes

**DOI:** 10.1101/2021.10.25.465765

**Authors:** Jacob M. Allen, Amy R. Mackos, Robert M. Jaggers, Patricia C. Brewster, Mikaela Webb, Chia-Hao Lin, Chris Ladaika, Ronald Davies, Peter White, Brett R. Loman, Michael T. Bailey

## Abstract

Psychological stress alters the gut microbiota and predisposes individuals to increased risk for enteric infections and chronic bowel conditions. Intestinal epithelial cells (IECs) are responsible for maintaining homeostatic interactions between the gut microbiota and its host. In this study, we hypothesized that disruption to colonic IECs is a key factor underlying stress-induced disturbances to intestinal homeostasis. Conventionally raised (CONV-R) and germ-free (GF) mice were exposed to a social disruption stressor (Str) to ascertain how stress modifies colonic IECs, the mucosal layer, and the gut microbiota. RNA sequencing of IECs isolated from CONV-R mice revealed a robust pro-inflammatory (*Saa1, Il18*), pro-oxidative (*Duox2, Nos2*), and antimicrobial (*Reg3b/g)* transcriptional profile as a result of Str. This response occurred concomitant to mucus layer thinning and signs of microbial translocation. In contrast to their CONV-R counterparts, IECs from GF mice or mice treated with broad spectrum antibiotics exhibited no detectable transcriptional changes in response to Str. Nevertheless, IECs from Str-exposed GF mice exhibited an altered response to *ex vivo* bacterial challenge (increased dual Oxidase-2 [*Duox2*] and nitric oxide synthase-2 (*Nos2*)), indicating that STR primes host IEC pro-oxidative responses. In CONV-R mice stress-induced increases in colonic *Duox2* and *Nos2* (ROS generating enzymes) strongly paralleled changes to microbiome composition and function, evidencing Str-mediated ROS production as a primary factor mediating gut-microbiota dysbiosis. In conclusion, a mouse model of social stress disrupts colonic epithelial and mucosal integrity, a response dependent on an intact microbiota and host stress signals. Together these preclinical findings may provide new insight into mechanisms of stress-associated bowel pathologies in humans.

## Introduction

Psychological stress is associated with disruptions to intestinal physiology. This is supported by studies in humans and animal models that have linked excessive levels of stress to increased risk of enteric infection and inflammatory bowel diseases and syndromes (1–4). The gut microbiota is a large and diverse community of microorganisms that continuously interact with host epithelial and immune components, a relationship that ultimately shapes numerous physiological responses during health and disease (5–8). Stress-induced modifications to the gut microbiota have been documented to occur in humans and across various types of animal models, underscoring a potential nexus between stress and intestinal health (9–11). However, the precise interactions that occur between microbe and host in response to stress are not fully elucidated.

Intestinal epithelial cells (IECs) consist of a diverse range of cells, including absorptive enterocytes, goblet cells, Paneth cells, enteroendocrine cells, and stem cells, that function together to maintain barrier integrity and coordinate proper immune responses to both pathogens and the endogenous microbiota (12). To perform these tasks, IECs have evolved an array of capabilities such as the production and release of mucins, antimicrobial peptides, inflammatory cytokines, and reactive oxygen species (ROS), which help maintain homeostatic immune responses while concurrently preventing overt microbial penetration into host tissue (6, 13, 14). Despite largely serving a protective role, emerging evidence indicates that abnormal epithelial responses to the endogenous microbiota may be a key component leading to gut dysbiosis and bowel disease manifestation (15). This may be in part due to chronic unabated stimulation of inflammatory pathways within IECs. For instance, hyperactivation of the NFκB pathway through aberrant toll-like receptor (TLR) or nod-like receptor (NLR) signaling on IECs is considered a major risk factor for inflammatory bowel diseases, such as Crohn’s disease and ulcerative colitis (16–18). Consistent with this role, we recently reported that an IEC-specific ablation of the classical NFκB pathway beneficially modified the gut microbiota and inhibited infectious colitis by *Citrobacter rodentium* in mice (19). This is in contrast to stress exposure, which alters the microbiota and exacerbates gastrointestinal inflammation associated with *C. rodentium* infection (20, 21). Ultimately, these data lead us to postulate that stressor-induced shifts in the gut microbiota and susceptibly to enteric colitis may be linked through alterations in epithelial physiology.

Herein, we utilize a well-validated social disruption stressor (Str) to investigate the effects of psychological stress on the colonic epithelial layer. Our results from conventionally-raised and germ-free animals support the hypothesis that stress induces a global shift in IEC gene transcription indicative of heightened pro-inflammatory, antimicrobial, and ROS signaling; an effect that is dependent on the gut microbiota. Concomitant disruption to the structural integrity of the mucosal layer and amplified systemic markers of bacterial translocation (LBP) signify Str-induced interferences to normal barrier integrity, a phenomenon through which IECs appear to play a central role. Lastly, we demonstrate that Str-induced augmentation in genes involved in IEC ROS production (*Duox2* and *Nos2*) paralleled changes to microbiota catalase activity, together indicating disruptions to IEC-gut microbiota communication with potential implications for gastrointestinal health.

## Results

### Social Disruption Stress Initiates a Unique Transcriptional Signature within IECs

We first exposed conventionally-raised (CONV-R) C57Bl/6 mice to a social disruption stressor (Str, 2 h/d) or a home cage control setting (Con) for 6 d. 12 h after the final Str exposure, colonic intestinal epithelial cells (IECs; EpCAM+, CD45-) were isolated and expression profiles analyzed by RNA sequencing. Using the differential gene expression analysis (DESeq2), we identified a broad upregulation in epithelial gene transcription induced by Str (Figure 1a; Volcano plot, 166 genes upregulated, 16 downregulated, Log Fold Difference > 0.5, Adj p <10^-5^). Over representation pathway analysis (ORA; NetworkAnalyst; (22)) using the Gene Ontology (GO:BP) database revealed numerous pathways affected by stress (FDR p<0.05; Top 10 enriched pathways shown in Figure 1b). This included Str-induced activation of immune (GO:BP *Inflammatory response;* genes represented in heatmap shown in Figure 1c) and antimicrobial (GO:BP *superoxide signaling* and *defense response*) pathways that occur concomitant to upregulation in pathways related to direct bacterial signaling (GO:BP *defense response to bacterium*, *NFκB cascade*; FDR p<0.05; Figure 1b). ORA analysis using the KEGG database revealed similar changes in pro-inflammatory and antimicrobial signaling pathways (**Figure S1a**), including a robust upregulation in genes associated with the NOD-like and Toll-like receptor (NLR, TLR) signaling pathways (FDR p<0.05; **Figure S1b-c).** In addition, IEC transcriptional profiles revealed a Str-induced downregulation in genes involved in fatty acid degradation and peroxisome biosynthesis that occurred alongside increased expression of genes involved in central carbon metabolism, together indicating coordinated metabolic shifts within IECs in response to Str (FDR p<0.05; **Figure S1d-e**). Further network visualizations using the GO:BP and KEGG pathways revealed Str-induced coordinated metabolic, hypoxia, wounding response and cytokine-mediated signaling pathways (among others) that corresponded to enhanced immune and antimicrobial signaling (FDR p<0.05; **Figure S2a-b**). Lastly, we confirmed that protein levels of the antimicrobial peptide “regenerating islet-derived protein 3-β” REG3β (encoded by *Reg3b*) reflected changes in IEC gene expression via immunofluorescence (p<0.05; Figure 1d).

**Figure 1.**
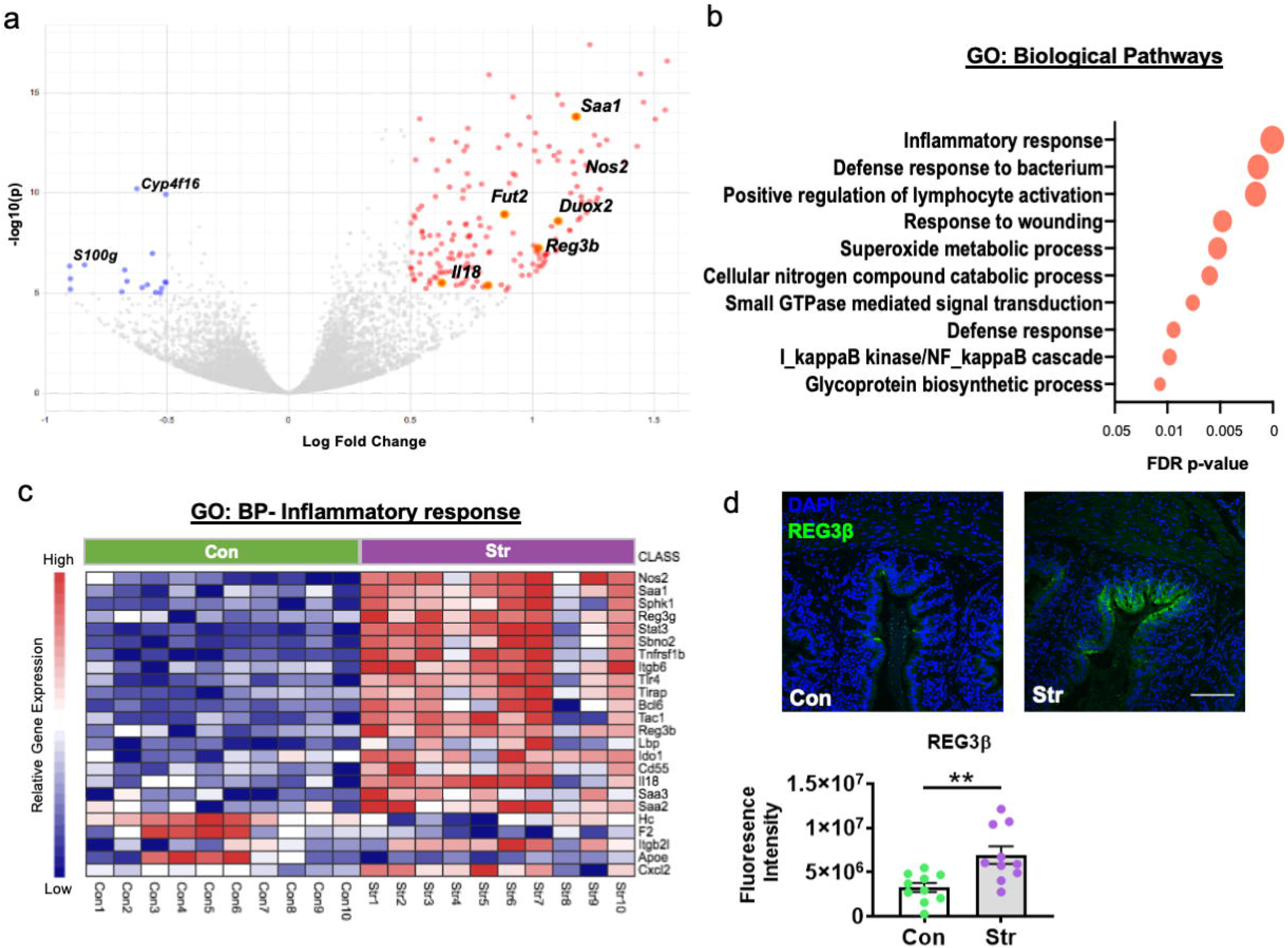
Stress shifts IEC transcription towards heightened inflammation, ROS production and antimicrobial defenses. **(a)** Volcano plot based on differential expression analysis (DESeq2) from RNA sequencing data reveals broad modifications to colonic IEC (EPCAM+, CD45-) gene expression in response to social stress (Con vs. Str [n = 10/group]; 166 genes upregulated by Str (red), 16 downregulated by Str (blue); Log Fold Change>0.5, Adjusted p<10^-5^. **(b)** Differential pathway analysis (NetworkAnalyst (23) of RNA sequencing data indicated Str-induced upregulation in pathways (Gene ontology: Biological Pathways [GO:BP]) related to immune, ROS and anti-microbial defense signaling, among others (Top ten shown; FDR p<0.05) **(c)** Heatmap of significantly altered genes (Con [green] vs. Str [purple]) within the GO:BP Inflammatory response pathway (Adjusted p<10^-5^) (d) Immunofluorescence staining confirms upregulation in protein levels of the antimicrobial peptide “regenerating islet-derived protein 3-beta” (REG3β), in the colon of Str-exposed mice, * = p<0.05, n = 10/group. Scale bar = 100 µM.

### Stress Disrupts the Colonic Mucus Layer Concomitant to Signs of Systemic Bacterial Translocation

We next explored the potential source of enhanced inflammatory and antimicrobial signaling observed in IECs. In the colon, a two-tiered mucus layer maintains a biophysical barrier that prevents excessive bacterial adhesion to IECs and abnormal infiltration into host tissue (23, 24). In addition to signs of microbial signaling, we also observed increased expression of genes involved in mucin/glycoprotein biosynthesis within stressor-exposed IECs (DESeq Adj p<10^-5^; Figure 2a). Therefore, we next asked whether structural changes to the colonic mucus layer may also occur as a result of stress. In a separate cohort of Str-exposed and Con mice (n =12/group), we used fluorescence in situ hybridization targeting endogenous bacteria (FISH; EUB338-Cy3) alongside lectin-based staining of the terminal mucus sugar, fucose (UEA1-fluorescein), to visualize potential changes to the integrity of the colonic mucus layer relative to the endogenous microbiota (Figure 2b). Using qualitative and quantitative scoring systems, we found that 6 d of Str induced a significant disruption to overall mucosal integrity alongside a concurrent reduction in inner mucus layer thickness (p<0.05; Figure 2c-d). While an intact layer with clear separation between the microbiota and IECs was observed in unstressed-control mice, Str-exposed mice exhibited a thinner, more porous and unstructured outer mucus layer with increased evidence of bacterial penetration into the inner mucosal layer (Figure 2b-d). This included evidence of bacterial adhesion to IECs in Str-exposed mice, which rarely occurred in Con mice (**Figure S3a**). Next, we explored whether Str initiated changes to goblet cell density (the primary mucus producing cell in the colon). However, no changes to total goblet cell number or density per crypt were observed in response to Str (**Figure S3b-c**). Nevertheless, alongside evidence of bacterial penetration within colonic tissue, Str-exposed mice exhibited higher levels of serum lipopolysaccharide-binding protein (LBP) compared to Con mice, indicating bacterial translocation occurs in parallel to colonic mucus disruption (p< 0.05; Figure 2e).

**Figure 2.**
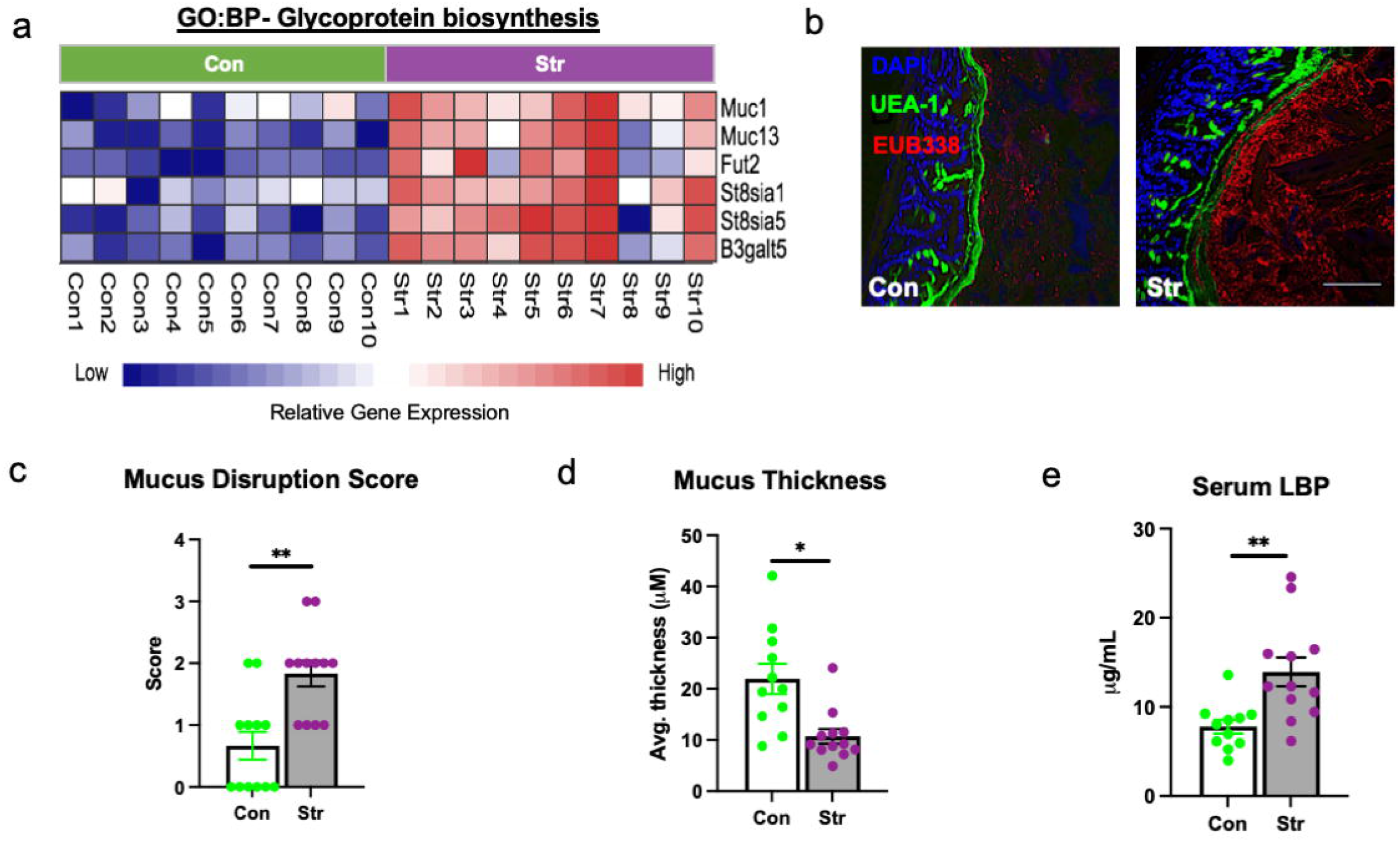
Stress disrupts the colonic mucus layer. **(a)** Heatmap of significantly altered IEC genes (Con [green] vs. Str [purple], n = 10/group) involved in glycoprotein biosynthesis as determined by RNA sequencing pathway analysis (GO:BP, DeSeq2 confirmed; p<10^-5^). **(b)** Representative FISH staining of total bacteria (EUB-338-Cy3), lectin-based fluorescence staining targeting terminal fucose sugars on mucins (UEA1-fluorescein; fucose) and host nuclei staining (DAPI) within colons of Con and Str-exposed mice**. (c)** Colonic mucus integrity score in response to Str (0 = normal mucus thickness with bacterial adherence only on outer mucus layer, 1 = normal mucus thickness and integrity with minor bacterial penetration to inner mucus layer, 2 = moderate outer mucus layer depletion with moderate bacterial penetration to inner mucus layer, minor evidence of bacterial adherence to epithelial layer, 3 = moderate depletion of outer and inner layer integrity with high bacterial penetration to inner mucus layer and moderate bacterial adherence to epithelium, 4 = major mucus depletion of outer and inner layer with substantial bacterial adherence to epithelium). **(d)** Inner mucus layer thickness (Avg thickness (µM) measured over 5 crypt regions). **(e)** Serum lipopolysaccharide-binding protein (LBP) concentrations (µg/mL). Scale bar = 100 µM, n = 12/group, * = p<0.05.

### Stress-Induced Changes to the Epithelial Transcriptome is Dependent on an Intact Microbiota

In light of combined evidence of altered mucus integrity and elevated signs of direct bacterial signaling within IECs, we next aimed to establish whether the presence of the gut microbiota was essential for mediating the IEC response to Str. First, we exposed a separate and independent cohort of CONV-R mice with an intact microbiota to the social defeat stressor (n=9/group). Using rt-PCR we confirmed that expression of a set of 6 genes (representative of the most altered pathways in the RNA-sequencing dataset: *Reg3b* (defense response to bacterium), *Duox2 and Nos2* (superoxide metabolic processes), *Fut2* (glycoprotein biosynthesis), *Saa1* and *Il18* (inflammatory response) were altered in IECs in response to Str (p<0.05; Figure 3a). However, when we exposed germ-free mice (GF; n=9/group) to the social defeat stressor, analysis of these same genes revealed an IEC transcriptional profile entirely unresponsive to the Str (p> 0.05 for all genes; Figure 3b). In addition to lack of differences in IEC expression profiles in response to Str, GF mice did not exhibit any changes to Str serum LBP compared to Con mice (p<0.05; **Figure S4**). GF mice have an altered mucosal layer and IEC physiology compared to mice that are born with an intact microbiota (CONV-R). This is highlighted by an overall lower expression of ROS (*Duox2* and *Nos2*) inflammatory (*Saa1*) and antimicrobial (Reg3b) genes in GF vs CONV-R mice (p<0.05; **Figure S4b**). In light of this distinctive physiology of GF mice, we deemed it necessary to also examine IEC transcriptional responses to stress in CONV-R mice exposed to a broad spectrum antibiotic cocktail (Abx) that significantly depletes microbial load in the distal colon. Similar to what was observed in GF mice, we did not observe a stress-induced upregulation of genes involved in ROS production, antimicrobial defense or innate inflammation in mice treated with Abx throughout the 6 ds of the Str paradigm (Figure 3c). In fact, in mice treated with Abx, IEC expression of *Saa1* and *Il18* were significantly reduced by Str, albeit through currently unknown mechanisms.

**Figure 3.**
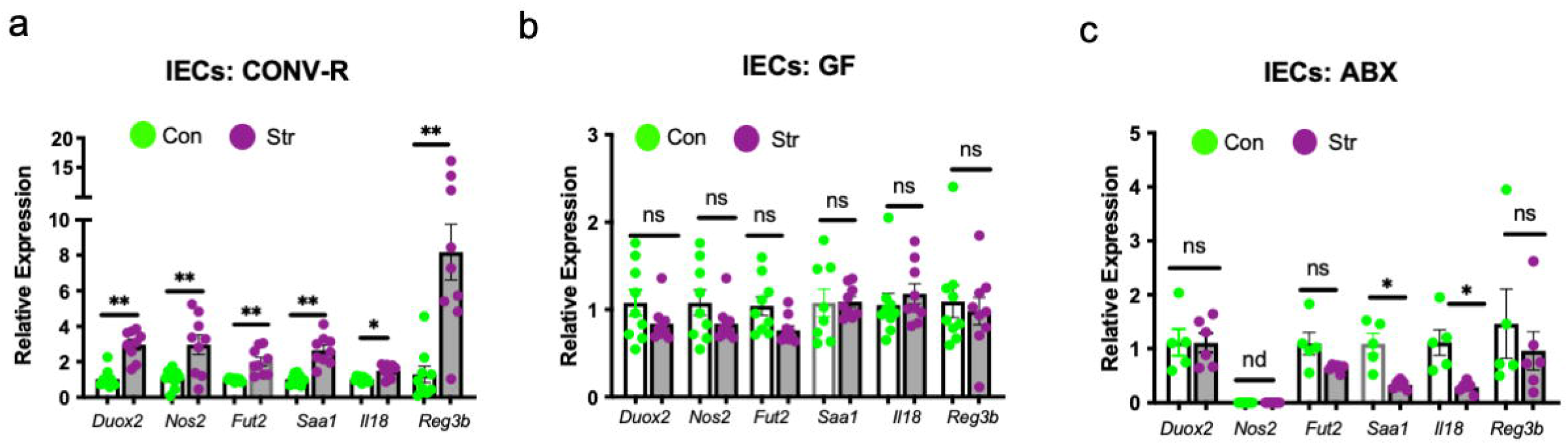
Stress-induced modifications to IECs are dependent on an intact gut microbiota. RT-PCR was used to analyze the expression of a select set of genes representing ROS-generating (*Duox2, Nos2*), mucus regulating (*Fut2*), innate inflammatory (*Saa1*, *Il18*) and antimicrobial (*Reg3b*,), signaling pathways within IECs (EpCAM+ CD45-) isolated from **(a) c**onventionally-raised **(b)** germ-free and **(c)** broad spectrum antibiotic treated C57Bl/6N mice exposed to Con or Str conditions. n = 6-9/group; ns = not significant at p>0.05, * = p<0.05, ** = p<0.01, *** = p<0.001. nd= not detected.

### Stress-Induced IEC Duox2 and Nos2 Relies on Microbial and Host Signals

Despite no transcriptional changes of key IEC stress-response genes in GF mice, we deemed it reasonable that endogenous host stress signals may still play a role in mediating IEC physiology, albeit in combination with microbial signals. To explore this hypothesis, we exposed another cohort of GF mice to an identical social disruption paradigm (Str) followed by IEC isolation and culture for 2 h. Cultured IECs were then subjected to an additional 2 h of immune challenge with bacterial-derived lipopolysaccharide (LPS) or flagellin (FLG) before RNA isolation and gene expression analysis. In analyzing a subset of 6 genes in IECS that were altered by Str in CONV-R, we found that dual oxidase 2 (*Duox2*) and nitric oxide synthase 2 (Nos2), two genes involved in ROS production, were primed by Str to over-respond to FLG and LPS challenges, respectively (Stress x LPS/FLG p<0.05, Figure 4a,). Stress exposure did not affect the response of other selected genes (*Fut2*, *Saa1*, *Il18, RegIIIb*) to an *ex vivo* challenges (Str x LPS/FLG, p > 0.05; Figure 4a, Table S1). We reasoned that a Str-induced increase in the expression of toll-like receptor-4 (TLR)-4 (the canonical host receptor for LPS) or toll-like receptor-5 (TLR)-5 (the canonical host receptor for FLG) may underlie the potentiation of *Duox2* or *Nos2* expression by Str. However, we failed to find any significant Str-induced changes to IEC expression of *Tlr4* or *Tlr5* at baseline or in response to an *ex vivo* bacterial challenge in GF mice (Str x LPS/FLG p>0.05; **Figure S5a**). To determine if these host-microbiota interactions were dependent on an in-tact microbiota *in-vivo*, we next used identical stress and *ex vivo* methods in CONV-R mice, focused on IEC expression of the same target genes described above. Unlike in GF mice, we did not observe any interactions of Str and *ex vivo* immune challenge on IEC gene expression in CONV-R mice. Nevertheless, we again observed strong main effects of Str on expression of *Duox2 and Nos2 (*as well as *Fut2, Saa1 and Reg3b)* across all *ex vivo* conditions (p<0.05; **Figure S5b** and Table S2). We deemed it possible that the difference in GF and CONV-R mice responses to FLG and LPS challenge may be related to Toll-like receptor expression in IECs. However, we failed to find any differences in IEC *Tlr4* or Tlr5 expression in GF vs CONV-R mice (**Figure S5c**). In light of the unique sensitivity of *Duox2* to Str and *ex vivo* challenges, we next investigated whether colonic DUOX2 protein mimicked gene expression changes. Indeed, immunofluorescence of colonic tissue revealed higher levels of colonic DUOX2 protein expression in Str mice, with notable fluorescence signaling observed on the luminal side of the epithelium (p<0.05; Figure 4b).

### Stress Modifies Gut Microbiota Composition Parallel to IEC Transcriptional Changes

Changes to epithelial physiology have been linked to gut microbiota dysbiosis (15, 25). Thus, we next explored the potential that stressor-induced augmentation in IEC genes (related to inflammation, and antimicrobial and ROS generation) would associate with changes to microbiota composition. To accomplish this, we implemented 16S rRNA gene sequencing to analyze the colonic microbiome of mice from our initial IEC RNA sequencing experiment. Using principle coordinate analysis (unweighted UniFrac), we confirmed the previous findings of our laboratory (10, 26) and others’ (27, 28) that stress exposure induces a broad and significant shift to colonic microbiome community structure (ADONIS p<0.05; Figure 5a). Further taxonomic analysis revealed numerus bacterial genera altered by stress, including a robust increase in relative abundance of *Muribaculaceae, Enterorhabdus, Marvinbryantia,* and *Candidatus Arthromitus* that occurred alongside a significant decrease in the abundance of genera from the *Lachnospiraceae* family (FDR p<0.05; Figure 5b). Out of these differentially altered bacteria, we noted that the abundance of *Muribaculaceae* (the most upregulated taxa) covered a wide range within Str-exposed mice (4-52% of total bacteria). In light of this variation, as well as the known communication between IECs and the microbiome, we returned to the RNA sequencing data to determine if the abundance of *Muribaculaceae* was related to IEC expression profiles. Among 182 differentially expressed genes in IECs (Con vs. Str, adj p < 10^-5^), we found that the expression of both *Duox2* and *Duoxa2* (a gene encoding the maturation protein for DUOX2), as well as *Nos2*, were associated with *Muribaculaceae* abundance (Spearman rho > 0.63, p<0.05; Top 20 differentially expressed IEC genes vs. *Muribaculaceae* shown in Figure 5c-d).

**Figure 4.**
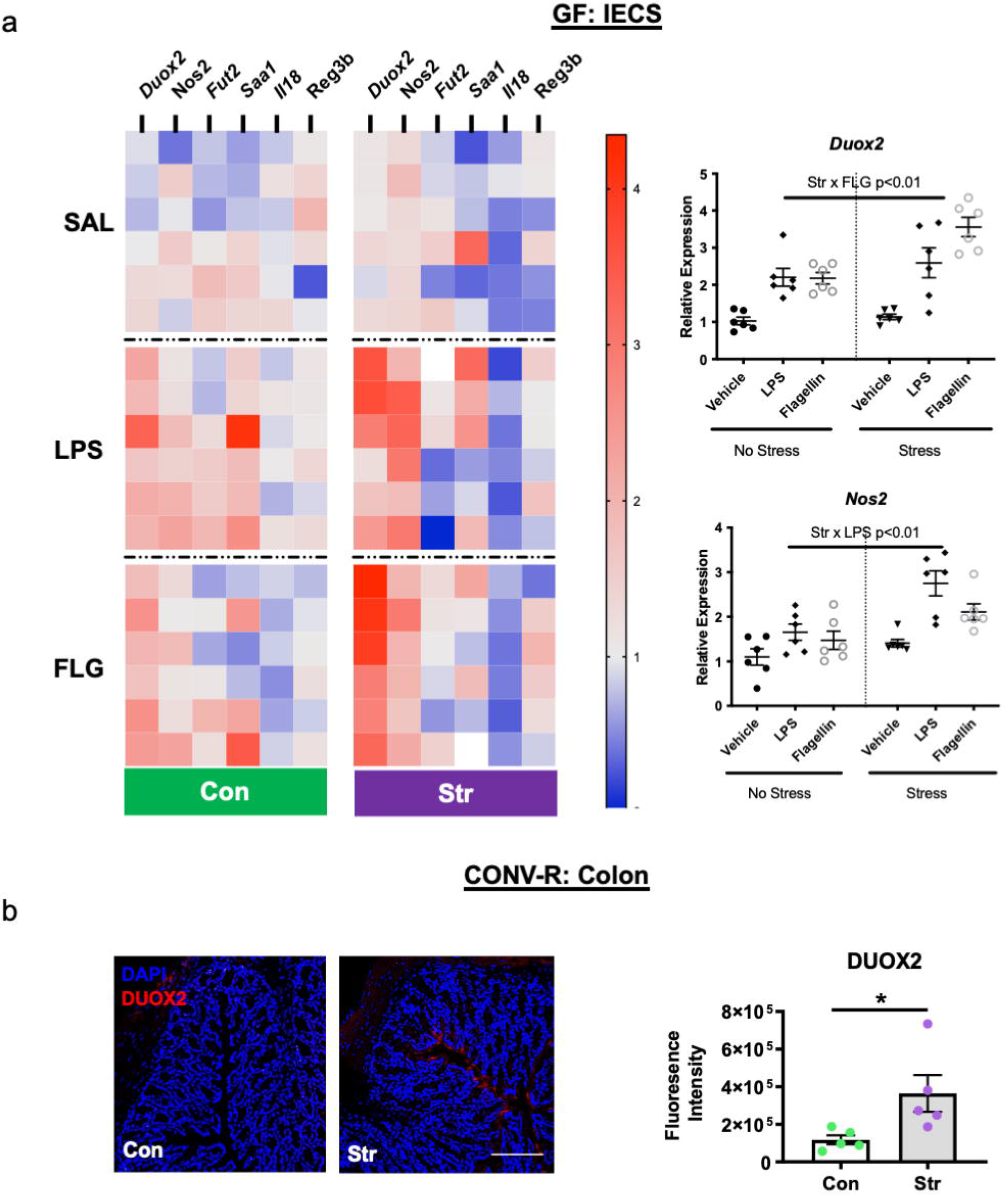
IECs are primed by host stress signals to respond to immunogenic bacterial components. IECs (EpCAM+, CD45-) isolated from germ-free (GF) Str-exposed and Con mice (n = 6/group) were cultured *ex vivo* before exposure to a vehicle (VEH-phosphate-buffered saline), lipopolysaccharide (E. coli LPS [1 μg/mL]) or flagellin (*S. typhimurium*-ultrapure FLG [200 ng/mL]) challenge for 2 h. **(a)** Heatmap representing relative gene expression of GF IECs exposed *in vivo* stress (Str) and *ex vivo* bacterial (LPS/FLG) challenge. Significant Str x LPS/FLG interactions (p<0.01) were found for *Duox2* and *Nos2*. For clarity, these data are re-represented in separate figures to the right of the heatmap. See Table S1 (GF) and Table S2 (CONV-R) for all means ± SEM and p-values for *ex vivo* experiments. **(b)** Representative images and quantification of DUOX2 protein expression from Con and Str conventionally-raised (CONV-R) mice (n= 6/group). Scale bar = 100 µM; * = p<0.05.

**Figure 5.**
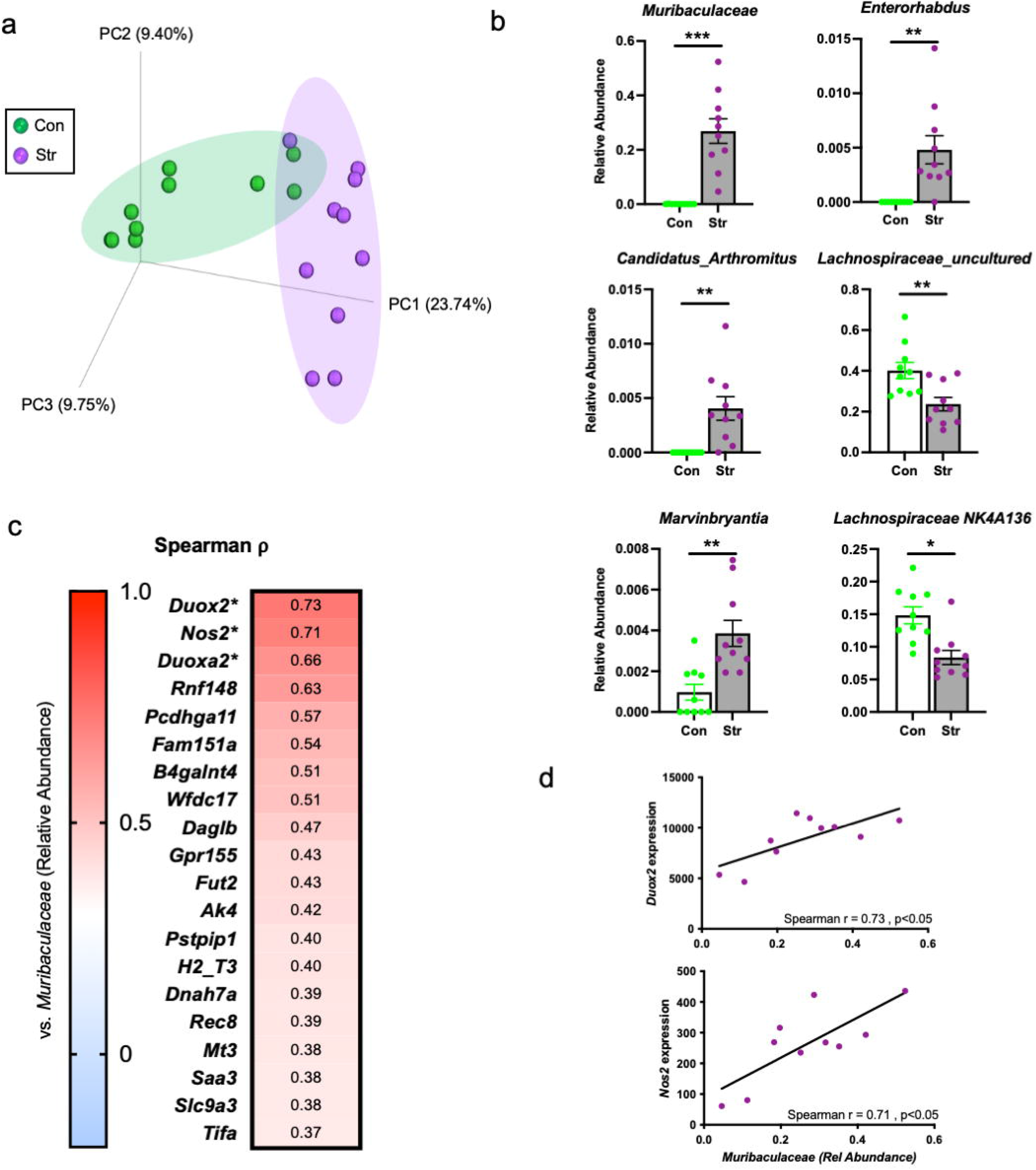
Stress modifies gut microbiota composition concomitant to changes in host epithelium. **(a)** Principle coordinates analysis based on the unweighted UniFrac distance metric reveals a robust shift in colonic microbiome composition in response to Str (ADNOIS p<0.05). **(b)** Bacterial genera significantly altered by Str exposure (top 6 altered differentially abundant genera shown). **(c)** Spearman rho (⍴) correlation coefficients relating *S24-7* abundance to IEC expression profiles (within Str group only; top 20 out of 183 differentially expressed genes shown). **(d)** *S24-7* relative abundance positively associates with IEC *Duox2* and *Nos2* expression (within Str group only). *** = FDR p<0.001, ** = FDR p<0.01, * = FDR p<0.05.

### Stress-Induced Modifications to Gut Microbiota Function and Activity Reflects Adaptations to IEC ROS Generation

DUOX2 produces hydrogen peroxide (H2O2), a ROS that, if produced at high levels, can be toxic to colonic anaerobic bacteria that do not contain the H2O2 degrading enzyme, catalase (29, 30). Thus, in light of the strong parallels between *Duox2* and *Duoxa2* expression and gut microbiome composition, we next aimed to establish whether Str (n=3/group) would shift the function of the microbiota towards a state of increased resistance to ROS production by IECs. Using metagenomic sequencing of distal colonic contents, we observed a Str-induced trend for increased overall catalase gene copy abundance, primarily mapping to *Bacteroides* and *Parabacteroides* genera (p=0.10; Figure 6a). Given this tendency, we next considered whether catalase enzyme activity would also increase as a result of Str. Indeed, in an independent cohort of mice exposed to the Str paradigm (n=5/group), we found augmented catalase activity within the colonic luminal contents of Str-exposed mice compared to Con mice (p<0.05; Figure 6b).

**Figure 6.**
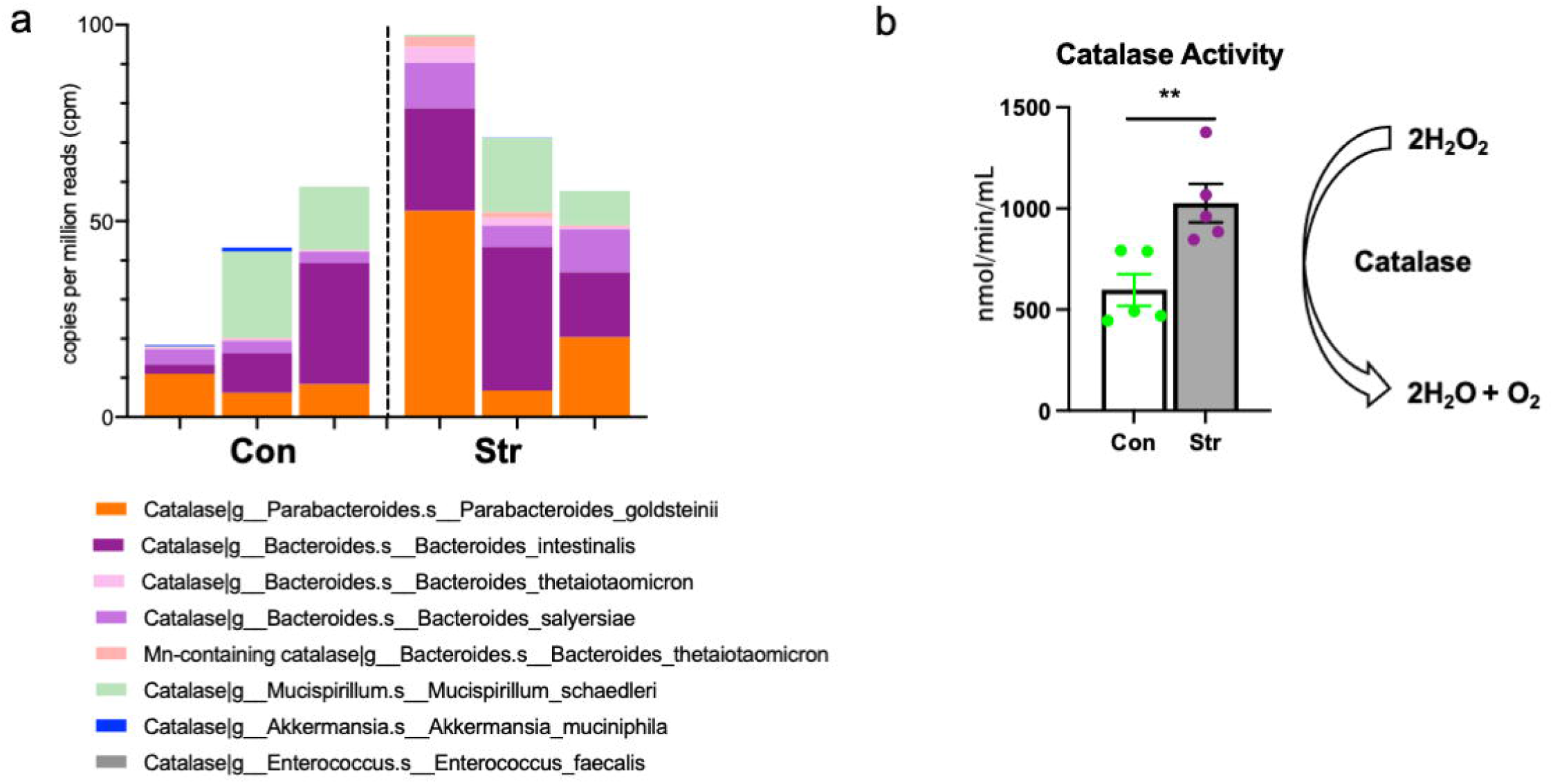
Stress modifies gut microbiota catalase gene abundance and enzyme activity. **(a)** Relative abundance of catalase gene content observed across bacterial taxa in colonic contents of Con and Str mice (n = 3/group). **(b)** Colonic luminal content catalase activity (nmol/min/mL) is upregulated in Str vs. Con mice (n = 5/group). * = p<0.05, ** = p<0.01.

## Discussion

Exposure to psychological stress alters the microbiota and predisposes individuals to increased risk of enteric infections and bowel diseases and conditions (1-3, 9, 20, 31, 32). However, stressor-induced modifications to the gut microbiota and gastrointestinal health have not been surveyed in context of intestinal epithelial cells (IECs), which have a unique role in: 1) maintaining a physical barrier between microbe and host, 2) relaying microbial signals to the immune system, and 3) producing bioactive molecules that subsequently modify the gut microbiota (33–35). In this study, we provide evidence that a mouse model of social defeat stress (Str) results in broad and physiologically relevant shifts to IEC physiology, underscored by an upregulation in antimicrobial, pro-inflammatory, and ROS-generating pathways that occur concomitantly to mucosal disruption, signs of systemic bacterial translocation, and functional changes to the gut microbiota.

Evidence of coordinated antimicrobial and immune signaling was present in Str-exposed IECs. This was apparent by transcriptional network analysis, which revealed an IEC gene signature supporting heightened innate immune signaling downstream of toll-like receptor (TLR) and nucleotide-binding oligomerization domain-like receptor (NLR) pathways. Constitutively expressed bacterial components such as LPS, FLG, and muramyl dipeptides bind TLRs and NLRs to activate the NFκB pathway, which ultimately enhances the expression of genes involved in innate immune signaling and ROS generation, such as *Saa1*, *Nos2* and *Duox2* (36). This enhancement in IEC TLR/NLR-to-NFκB signaling likely has important implications for understanding how stress predisposes to enteric infection and inflammatory bowel disease. Indeed, studies have demonstrated that hyperactivation of the NFκB pathway contributes to aberrant inflammatory responses to enteric infection and colitis (37, 38). In addition, recent data from our laboratory showed that ablation of the classical NFκB pathway in IECs conferred increased host protection against a colonic *Citrobacter rodentium* infection challenge (19). This is in contrast to stress, which exacerbates *C. rodentium* infection (3, 20, 21). Further studies are needed to unravel the mechanistic underpinnings of IEC NFκB signaling in response to stress and its potential role in mediating gastrointestinal infection and/or inflammatory bowel disease.

Under homeostatic conditions, the colon relies on two mucus layers to provide biochemical and physical protection against aberrant microbial signaling (23). We found that stress reduced the thickness and disrupted the integrity of these mucus layers. Morphological changes to the mucus coincided with altered transcription of genes involved in glycoprotein biosynthesis (e.g. *Muc1*, *Muc13*) and mucin glycosylation (e.g. *Fut2*, *St8sia1*) in IECs. While mechanisms underlying stress-induced mucus disruption remain unknown, it is important to note these transcriptional and morphological changes did not occur in GF mice, highlighting a key role of an intact microbiota in modifying the mucus layer during Str. Indeed, other research has identified specific members of the microbiota, such as *Bacteroides thetaiotamicron*, that are directly responsible for modifying mucus integrity and barrier function (39, 40), some of which are associated with mucus disruption during inflammatory bowel diseases like ulcerative colitis and Crohn’s disease (40, 41). In this study, it is feasible that *Muribaculaceae*, which was highly upregulated by stress and was recently found to exhibit robust mucus degrading properties (42), may be driving changes to mucus integrity and bacterial translocation observed in response to stress. Indeed, our data coincide with previous reports of social stress-induced bacterial translocation (20, 43) and highlight the need for further research into the role of colonic mucosal integrity as a mechanism by which endogenous microbes (or immune-stimulating bacterial components) reach host circulation during stress. Future studies using gnotobiotic models (monocolonization or multi-strain colonization) will be needed to help ascertain the specific microbes responsible for stress-induced IEC transcriptional responses, mucosal layer degradation and barrier disruption.

It is likely that exogenous signals, in addition to direct microbial stimulation, contribute to increased activity of IECs. This includes Reg3b and Reg3g, antimicrobial C-type lectins whose transcription activity within IECs are strongly upregulated by exogenous signals (44, 45). While not a focus of this study, cytokines released in response to bacterial signaling by local immune cells (e.g. innate lymphoid cells & Th17 cells) can activate anti-microbial defense pathways in epithelial cells (46, 47). In support of this, our transcriptome data revealed Str-induced gene transcription profiles centered around activation of the Janus kinase/Signal transducer of activated transcription factor-3 (JAK/STAT3) pathway, which is activated downstream of IL-22 and IL-17 (among other cytokines) and is the primary route of *Reg3b* and *Reg3g* transcription in IECs (48). The specific effects of stress on the colonic immune cell niche, however, remains understudied. Nevertheless, our network analysis of IEC expression profiles revealed Str-induced upregulation in genes involved in Th17 differentiation (e.g. *Saa1/3*), highlighting a potential role of IECs in driving local changes to the innate and adaptive immune cell compartments, which may ultimately feedback to control IEC physiology (35). Future studies will need to delineate the mechanisms and implications of an immune-epithelial cell axis in response to stress.

In germ-free mice IEC *Duox2* and *Nos2* expression (while unchanged at baseline) were uniquely primed by host stress signals to respond to bacterial (Flagellin and LPS) challenge. *Duox2* encodes DUOX2 protein, which is a member of the NAD(P)-H dependent oxidase family that is highly expressed on IECs and is the primary contributor of host hydrogen peroxide (H2O2) production in the mammalian GI tract (49, 50). *Nos2* encodes inducible nitric oxide synthase (iNOS), an enzyme that produces nitric oxide (NO) in response to inflammatory signals. NO is a relatively weak free radical, but contributes to toxicity in the gut by forming peroxynitrite (NO3^-^) (51, 52). DUOX2 and iNOS mediated ROS and RNS production represent important anti-bacterial defense pathways within the mammalian colon (15, 53). For example, DUOX2-mediated ROS generation was shown to be required for an adequate NOD2-mediated antimicrobial response to *Listeria monocytogenes* challenge (25). Meanwhile, iNOS mediated NO production limits growth of enteric pathogens such as *Salmonella Tymphimurium* (54). However, while acute DUOX2 and iNOS signaling appears important for antimicrobial defense to enteric pathogens, unabated and chronic ROS signaling may cause long-term disruption to the gut microbiota and the epithelial niche. This is supported by a growing body of literature linking chronic DUOX2 and iNOS signaling to inflammatory bowel disease (IBD) and microbial dysbiosis (53, 55, 56). For example, both colonic DUOX2 and iNOS are upregulated during the early onset IBD (57, 58) while DUOX2 is highly responsive to a dysbiosis microbiome isolated from IBD patients (15). Future studies that delineate the role of stress-induced modifications to DUOX2 and iNOS activity may help unravel the complex etiology of stress-associated inflammatory bowel diseases.

Catalase, an enzyme that speeds the breakdown of H2O2 into oxygen and water, represents a main line of bacterial defense against host-mediated ROS generation (59). However, catalase is not expressed by many commensal anaerobic microbes that reside in the colonic niche, leaving them vulnerable to excess ROS production (60, 61). In the case of stress, we hypothesized that high production of H2O2 through enhanced DUOX2 signaling may deplete catalase deficient microbiota species. In line with this premise, we found that each taxa reduced by Str, including bacteria from *Lachnospiracae* family, are almost entirely catalase negative. Conversely, species within the most prominently upregulated taxa in response to stress, species from the family *Muribaculaceae* (previously named S24-7), expresses catalase (62). Based off of these findings, it is tempting to speculate that catalase-positive bacteria are able to survive Str-induced upregulation in DUOX2 activity and subsequent H2O2 production, whereas bacteria without the enzyme are depleted. Indeed our data showing increased catalase enzyme activity in the colonic lumen of Str-exposed mice support the idea that catalase-positive bacteria, regardless of taxonomy, bloom in response to Str. In fact, previous work in mice (26) and in humans (4, 9) has shown that stress exposure is associated with increased abundance of *Bacteroides spp.* and *Parabacteroides spp*., oxygen tolerant anaerobic commensals found in the mammalian microbiome, both of which express catalase (30). In addition, IEC DUOX2 activity can promote a competitive niche for catalase-positive facultative anaerobes such as *E. coli* and *C. rodentium,* while inducing a hostile environment for commensals such as *Lactobacillus spp.* (14). This is consistent with stress-induced reductions in *Lactobacillus spp*. abundance and increased *C. rodentium* colonization (3, 63, 64). Future studies are needed to further unravel how the microbiota and pathogens respond to enhanced ROS-generation by IECs during stress.

The effects of psychological stress on the mammalian gut and its microbiota are only beginning to be explored. In this study, we show that a mouse model of psychological social stress induces a robust and reproducible transcriptional signature within IECs indicative of enhanced pro-inflammatory, antimicrobial, and ROS signaling. These effects, which were reliant on an intact microbiota, manifested in morphological changes to the mucus layer and signs of bacterial translocation, underscoring the significance of the intestinal epithelial layer as a central component to stress physiology. We also uncovered a potential role of epithelial cells as mediators of gut microbiota function, evidenced by Str-induced increase in microbe-specific ROS detoxifying activity (catalase) that paralleled ROS-generating capacity of IECs. Together, our data uncover a maladaptation to psychosocial stress involving the gut microbiota and intestinal epithelial cells that provides a strong link to enteric infection and bowel disease manifestation.

## Materials and Methods

### Animals

Adult male C57BL/6NCrl strains of *Mus musculus* (Charles Rivers Laboratories, Wilmington, MA) between 6-8 weeks of age were used in the study. Animals were given *ad libitum* access to water and autoclaved chow throughout all experiments (12 h light cycle; lights on at 06:00). Separate cohorts of CONV-R mice were used in the following experiments: 1. IEC RNA sequencing (n = 10/group), 2. Mucosal integrity (n = 9/group), 3. DUOX2 IF staining (n = 6/group) 4. Metagenomics (n = 3/group) and 5. Catalase activity (n = 5/group). 6. broad spectrum antibiotics (n=6/group). Germ-free mice (4 wks old) were purchased from Charles River. To prevent microbial contamination germ-free animals were kept in Sentry sealed positive pressure cages (Allentown, Allentown, NJ) for a 2-week acclimation period and over the course of the study. Germ free sterility was confirmed by following the standard contamination protocol used by the National Gnotobiotic Rodent Resource Center (NGRRC) at UNC-Chapel Hill (UNC). A separate cohort of GF mice were used for the following experiments: 1. IEC gene expression (n = 9/group) and 2. IEC *ex vivo* challenge (n = 6/group), totaling two separate cohorts of GF mice. All experimental procedures were conducted in the Animal Resources Core (ARC) facility at Research Institute at Nationwide Children’s Hospital or at the Animal Care Facility at the University of Illinois at Urbana-Champaign with the approval of the Research Institutional Animal Care and Use Committees at each institution (NCH-Protocol #AR15-00046 and UIUC-Protocol # 20241).

### Social disruption stressor

Social disruption stress (Str) was completed by placing an aggressor mouse (retired breeder C57BL/6NCrl) into the cage with 3 resident mice of the same strain for 2 h between 16:30-18:30 occurring at the end of the dark cycle. Str was deemed effective when resident mice exhibit subordinate behaviors as described previously (26). This protocol was repeated for 6 consecutive d with animal sacrifice and tissue collection occurring 12-15 h after the final stressor. These mice were closely monitored, but none of the mice needed to be removed for excessive wounding using criteria previously described (22). Mice in the control condition were not handled throughout the experiment. GF aggressors were checked for microbial contamination by PCR and culture in an identical manner to experimental mice

### Antibiotic Treatment

A broad-spectrum antibiotic cocktail (∼40mg/kg/day, ampicillin 33 mg/kg/day, metronidazole 21.5mg/kg/day, vancomycin 4.5mg/kg/day; Sigma-Aldrich) was provided in drinking water throughout the 6ds of the Str paradigm.

### Intestinal epithelial cell isolation and culture

Colons were removed from animals immediately after sacrifice, opened longitudinally, washed with PBS thoroughly and cut into 5 mm pieces. Tissue was then placed in a pre-digestion solution (1x HBSS, 5 mM EDTA, 1mM DTT, 5% FBS with Antibiotic/Antimycotic solution (Sigma Aldrich, St. Louis, MO)) and rotated for 20 min at 37°C. After brief vortexing at high speed (10 s), the tissue homogenate was filtered with a 100 µM mesh filter with the resulting pass through (containing IEC fraction) placed on ice. The remaining colon pieces were placed in 20 mL of fresh pre-digestion buffer and rotated again. These steps were repeated 3 consecutive times with pass through stored on ice after each rotation to ensure adequate IEC removal from lamina propria. Next, single cell suspensions containing the IEC fraction were diluted to 10^8^ cells/mL in MACs buffer (0.5% BSA 2mM EDTA) before being processed with the Miltenyi Dead Cell Removal Kit (Miltenyi Biotec, Auburn, CA) per manufacturer’s instructions. After a wash step, live cells were incubated with CD45 magnetic beads for 10 min before being passed through Miltenyi LS columns per manufacturer’s instructions. After an additional wash step, the CD45^-^ cell fraction (10^7^-10^8^ cells) was incubated with EpCAM^+^ beads for 10 min before again passing through LS columns per manufacturer instructions. Resulting cells were collected for RNA isolation. For *ex vivo* experiments, cells were plated in 24-well cell culture plates at 2 x 10^5^ cells/well and incubated (5% CO2) at 37°C for 2 h. Cells were then treated with saline (PBS), LPS (*E. coli* O55:B55 (1 µg/mL); Sigma Aldrich) or FLG (Ultrapure-*Salmonella typhimurium* (200ng/mL); Invitrogen, San Diego, CA) and incubated at 37°C for another 2 h before RNA isolation.

### RNA sequencing

IEC RNA was isolated using the PureLink RNA Mini Kit (Thermo Fisher Scientific, Waltham, MA) according to manufacturer’s instructions. Following assessment of the quality of total RNA using Agilent 2100 bioanalyzer and RNA Nano Chip kit (Agilent Technologies, Santa Clara, CA), rRNA was removed from 2000 ng of total RNA with Ribo-Zero rRNA removal kit specific for Human/Mouse/Rat. To generate directional signal in RNA seq data, libraries were constructed from first strand cDNA using ScriptSeq^TM^ v2 RNA-Seq library preparation kit (Epicentre Biotechnologies, Madison, WI). Briefly, 50 ng of rRNA-depleted RNA was fragmented and reverse transcribed using random primers containing a 5’ tagging sequence, followed by 3’end tagging with a terminal-tagging oligo to yield di-tagged, single-stranded cDNA. Following purification by a magnetic-bead based approach, the di-tagged cDNA was amplified by limit-cycle PCR using primer pairs that anneal to tagging sequences and add adaptor sequences required for sequencing cluster generation. Amplified RNA-seq libraries were purified using AMPure XP System (Beckman Coulter, Brea, CA). Quality of libraries were determined via Agilent 2200 TapeStation using High Sensitivity D1000 tape and quantified using Kappa SYBR®Fast qPCR kit (KAPA Biosystems, Inc, Wilmington, MA). Approximately 60-80 million paired-end 150 np sequence reads were generated per sample using the Ilumina HiSeq 4000 sequencer.

### RNA-seq data analysis

Each sample was aligned to the GRCm38.p4 assembly of the mouse reference from NCBI using version of 2.5.1b of the RNA-Seq aligner STAR (http://bioinformatics.oxfordjournals.org/content/29/1/15). Transcript features were identified from the general feature format (GFF) file that came with the assembly from the NCBI. Feature coverage counts were calculated using HTSeq (http://www-huber.embl.de/users/anders/HTSeq/doc/count.html). The raw RNA-Seq gene expression data was normalized, and post-alignment statistical analyses and figure generation were performed using DESeq2 (http://genomebiology.com/2014/15/12/550) and analyzed by the open-source NetworkAnalyst 3 software (22). Variance and low abundance filters were set at 15 and 4, respectively. Samples were normalized to Log2-counts per million reads. Normalized read counts were then analyzed (Con vs. Str) by differential expression analysis based on the binomial distribution (DESeq2 (65)) with significance set at Log2 Fold Change > 0.5 and an Adj p< 10^-5^. Over representation pathway (ORA) heatmap clustering, and network analysis was performed through NetworkAnalyst (22) with significantly altered genes (DeSeq2 Adj p<10^-5^) as input variables. Differences in pathway abundance were deemed significant at FDR p<0.05 through NetworkAnalyst.

### Collection and processing for immunofluorescence and FISH staining

Immediately after sacrifice, sections of the mid-colon were collected (with intestinal contents left in place) and tissues were fixed by immersion in methanol Carnoy’s solution (60% methanol, 30% chloroform, 10% acetic acid) for 48-96 h, followed by two successive washes each in methanol for 35 min, ethanol for 30 min, and xylene for 25 min. Cassettes were then submerged in melted paraffin at 68°C for 1 h, removed, and kept at RT until sectioning. Paraffin blocks were cut into 4 µm-thick sections and deparaffinized prior to immunofluorescence or FISH/mucus staining. For cryosectioning (DUOX2 staining), samples were snap-frozen in Tissue-Tek O.C.T. compound (Andwin Scientific, Woodland Hills, CA) and immediately sectioned at 8 µm thickness on a cryostat.

### Immunofluorescence

Sections were deparaffinized in 100% xylene for 10 min and 50% xylene/50% ethanol for 5 min followed by successive 5 min washes in 100%, 95%, 75%, 50%, 25% and 0% EtOH/PBS solution. Slides were then placed in citrate buffer antigen retrieval solution (pH 6.0) with 0.5 % Tween-20 in a 90°C water bath for 20 min. After allowing slides to cool in running H2O, slides were dried and incubated in permeabilization buffer (0.3% Triton-X, 1% BSA/PBS) for 30 min at room temperature followed by another 3 wash steps with PBS. Slides were then incubated in permeabilization buffer containing the primary REG3β antibody (1:250 dilution; R&D systems, Minneapolis, MN) overnight (16 h). After 3 consecutive washes, the slides were incubated in permeabilization buffer containing secondary antibody, anti-rabbit IgG-AlexaFluor 488 (1:500 dilution; Thermo Fisher Scientific), for 1 h in the dark. After an additional 3 washes (PBS), slides were dried and incubated with Prolong Gold Anti-Fade DAPI solution (Thermo Fisher Scientific) for 2 h in the dark prior to slide curation and imaging.

DUOX2 staining was completed on frozen sections as previously described (15). Briefly, thawed 8 µm sections were briefly fixed in 4% freshly prepared formaldehyde for 5 min, washed twice in PBS, and then blocked with 20% donkey serum in PBS. Primary pan-DUOX antiserum (1:1000; provided by Xavier De Deken’s laboratory) or normal rabbit IgG (control; Santa Cruz Biotechnology, Santa Cruz, CA) were applied to sections overnight at RT with slow rotation. After 3 consecutive washes anti-rabbit Alexa Fluor-568 conjugated antibodies (1:500; Thermo Fisher Scientific) were applied for 2 h before counterstaining with DAPI.

## FISH and mucus staining

Slides were deparaffinized by an initial incubation of slides at 60°C for 10 min and then incubated in xylene substitute solution twice for 10 min with the first solution pre-warmed to 60°C. Slides were then washed in 99.5% ethanol for 5 min and left to dry. For hybridization, slides were incubated in a pre-warmed hybridization solution (5M NaCL 1M Tris-Hcl (pH 7.4), 1% sodium dodecyl sulfate and 10% formamide) containing the FISH probe (EUB338-Cy3; see Table S3) before being placed in a sealed humid chamber at 49°C overnight. Slides were incubated with FISH washing buffer (0.9 M NaCl, 20 mM Tris-HCl (pH 7.4)) preheated to 49°C for 10 min, and washed three times in PBS. For mucus (fucose) staining, slides were then incubated with 1:500 fluorescein labeled Ulex Europaeus Agglutinin-1 (UEA-1-fluorescein,Vector Labs, Burlingame, CA) for 1 h in the dark. After 3 washes, slides were incubated with Prolong Gold Anti-Fade DAPI solution (Thermo Fisher Scientific) for at least 2 h in the dark prior to slide curation and imaging.

### Serum LPS-binding Protein

Serum LPS-binding protein concentrations were determined by a commercially available ELISA kit via manufacturer’s instructions (Hycult Biotec, Uden, Netherlands).

### Semi-quantitative Real-Time PCR

IEC RNA was isolated using the PureLink RNA Mini Kit (Thermo Fisher Scientific) according to manufacturer’s instructions. Complementary DNA was synthesized with a reverse transcriptase kit (Thermo Fisher Scientific). Power SYBR Green Master Mix (Thermo Fisher Scientific) with primers (Fwd and Rv, Integrated DNA Technologies, Coralville, IA) listed in Table S3 used together in PCR reactions. Differences in gene expression were determined by Real-Time PCR (Quantstudio 5, Thermo Fisher Scientific). Relative expression was determined using the delta-delta cycle threshold method (dd*C*t).

### 16S rRNA Gene Sequencing and Analysis

Intestinal contents were collected directly from the colon immediately after sacrifice and frozen at −80°C. Contents were homogenized and DNA was extracted (∼10 mg) using a QIAamp Fast DNA Stool Mini Kit (Qiagen, Hilden, Germany) following manufacturer’s instructions, with slight modifications. Contents were incubated for 45 min at 37°C in lysozyme buffer (22 mg/ml lysozyme, 20 mM TrisHCl, 2 mM EDTA, 1.2% Triton-x, pH 8.0), before homogenization for 150 s with 0.1 mm zirconia beads. Samples were incubated at 95°C for 5 min with InhibitEX Buffer, then incubated at 70°C for 10 min with Proteinase K and Buffer AL. Following this step, the QIAamp Fast DNA Stool Mini Kit isolation protocol was followed, beginning with the ethanol step. DNA was quantified with the Qubit 2.0 Fluorometer (Life Technologies, Carlsbad, CA) using the dsDNA Broad Range Assay Kit. Amplified PCR libraries were sequenced from both ends of the 250 nt region of the V3-V4 16S rRNA hypervariable region using an Illumina MiSeq. DADA2 and Quantitative Insights into Microbial Ecology [QIIME] 2.0 (66) were utilized for amplicon processing, quality control and downstream taxonomic assignment using the ribosomal RNA database SILVA version 138 (67). The EMPeror software package was used for three dimensional principle coordinates plotting of microbiome data (β-diversity; Unweighted Unifrac) (68).

### Metagenomic Sequencing and Analysis

DNA for metagenomic sequencing was extracted from 6 colonic content samples from a separate cohort of mice exposed to the stress paradigm (n=3/group) and sequenced at the Genomics Shared Research facility at The Ohio State University (Columbus, OH). Prior to library generation, microbial DNA enrichment was performed using the NEBNext Microbiome DNA Enrichment kit (New England BioLabs, Ipswich, MA) per manufacturers protocol. Libraries were generated using the NExtera XT Library System in accordance with the manufacturer’s instructions. Genomic DNA was sheared by sonication and fragments were end-repaired. Sequencing adapters were ligated, and library fragments were amplified with five cycles of PCR before solid-phase reversible immobilization size selection, library quantification, and validation. Libraries were sequenced on the Illumina HiSeq 4000 platform. All raw reads were trimmed from both the 5’ and 3’ ends with Sickle before downstream processing. 150 bp reads were processed downstream by HUMAnN 2.0. Briefly, the HUMANn2 core function was run using nucleotide mapping and translated search against the UniRef 90 database. To account for differences in sequencing depth, the resulting organism specific gene and pathway abundances were normalized to total reads (copies/million). For this analysis, relative catalase levels were tabulated on a per-taxa and total gene abundance basis and then analyzed across Str groups.

### Catalase Activity Assay

Colonic content samples that were flash frozen after sacrifice were thawed on ice and weighed to ∼100 mg. 1 mL of PBS was precisely added to every 100 mg of stool before being broken up by vigorous pipetting. Samples were vortexed for 1 minute and then spun at 10,000 x g for 5 min. To disrupt bacterial cell walls, contents were incubated for 45 min at 37°C in lysozyme buffer (22 mg/ml lysozyme, 20 mM TrisHCl, 2 mM EDTA, 1.2% Triton-x, pH 8) for 45 min at 37°C. Next, 300 mg of autoclaved beads (0.1 mm zirconia) were added to each sample before bead beating for 3 min. Samples were then spun down at 12,000 x g for 15 min. Supernatant was removed and stored on ice until catalase assay was performed. Catalase activity was determined using a fluorometric-based catalase activity kit according to manufactures instructions (Abcam, Cambridge, MA). Catalase activity was calculated as nmol/min/mL.

### Image Quantification

Immunostained and/or lectin stained cross-sectioned colon samples were analyzed by a LSM 880 confocal microscope (Zeiss Microscopy, Jena, Germany). After image capture, sections were analyzed for protein quantification through fluorescence intensity analysis in ImageJ. Mucus integrity was analyzed on FISH/mucus stained sections by a blinded observer with the following scoring system: 0 **=** normal mucus thickness with bacterial adherence only on outer mucus layer, 1 = normal mucus thickness and integrity with minor bacterial penetration to inner mucus layer, 2 = moderate outer mucus layer depletion with moderate bacterial penetration to inner mucus layer, 3 = moderate depletion of outer and inner layer integrity with high bacterial penetration to inner mucus layer and moderate adherence to epithelium, 4 = major mucus depletion of outer and inner layer with substantial bacterial adherence to epithelium. Inner layer mucus thickness was analyzed by ImageJ software. Goblet cells were counted and analyzed by total number and density per crypt. All measurements were made by a blinded observer and averaged across 5 randomly chosen crypt regions.

### Statistical Analyses

Student t-tests were used to analyze differences between Con and Str groups. Two-factor ANOVA with Str *(in vivo*) and LPS/FLG (*ex vivo*) as variables was used to analyze data from *ex vivo* IEC experiments. Tukey’s post-hoc was used for multiple comparison testing. Differences in microbiome β-diversity was determined through UniFrac distance metric (69) and analyzed by permutational multivariate analysis (ADONIS). Spearman rho non-parametric analysis was implemented to analyze associations between IEC expression profiles and the abundance of *Muribaculaceae*. The Benjamini-Hochberg false discovery rate (FDR) method was implemented to avoid type-1 error in all cases with multiple testing. Data is expressed as mean +/- SEM. Statistical alpha was set *a priori* to 0.05 for all analyses.

### Randomization and Blinding

ARRIVE guidelines for preclinical animal experiments were followed (70). All mice were randomly assigned to Str groups prior to the initiation of each experiment. Blinded observers were utilized for all staining quantification and histological scoring.

### Ethics Statement

All animals used in this study were approved by the IACUC at Nationwide Children’s Hospital. The authors report no competing interests.

## Data Availability Statement

The datasets used and/or analyzed during the current study are available from the corresponding authors on reasonable request. RNA-sequencing and 16S rRNA gene data will be deposited on NCBI.

## Acknowledgements

The authors thank X. De Deken, F. Minot and H. Grasberger for providing DUOX antibodies.

## CONTRIBUTORS

J.M.A. planned and performed experiments, collected and analyzed data, and wrote the manuscript. A.R.M. planned and performed experiments, collected data, and edited the manuscript R.M.J. P.C.B, C.H.L, M.W. and C.L performed experiments and edited the manuscript. B.R.L. and R.D analyzed data and edited the manuscript. P.W. collected data and edited the manuscript. M.T.B. planned experiments, analyzed data and edited the manuscript.

## List of abbreviations

IEC: Intestinal epithelial cell
ROS: Reactive oxygen species
CONV-R: Conventionally-raised
GF: Germ-free
*Il*: Interleukin
*Saa*: Serum amyloid a
*Duox*: Dual oxidase
*Reg3*: Regenerating islet-derived protein 3
*Nos2*: nitric oxide synthase-2

## Supplementary Figure Legends

**Figure S1.** Over enrichment (ORA) heatmap clustering analysis of IEC (EpCAM+CD45-) RNA sequencing data using using the KEGG database. **(a)** Top 10 differentially expressed pathways. **(b-e)** Heatmaps representing relative gene expression within selected pathways (KEGG) found to be significantly different between Con [green] vs. Str [purple], FDR p<0.05 by ORA heatmap clustering analysis.

**Figure S2.** ORA network analysis of IEC (CD45-, EpCAM+) RNA sequencing data. Genes entered into analysis were differentially expressed (Con vs. Str) based on the DESeq P Adj p<0.05. Pathways in network were derived through **(a)** KEGG and **(b)** GO:BP databases and were significantly different (Con vs. Str) at FDR p<0.05.

**Figure S3. (a)** Representative image displaying evidence of bacterial (EUB-338) penetration into mucus layer (UEA-1) and attachment to host epithelial cells (DAPI) in the colon of Str-exposed mice (integrity score = 2; arrows indicate signs of bacterial penetration into mucus layer [top arrow] and adhesion to host epithelial cells [bottom arrow]); scale bar = 20µM. **(b)** Goblet cell number and **(c)** density did not differ as a result of Str. * = p<0.05, ns = not significant.

**Figure S4. (a)** Stress does not alter serum LPS-binding protein (LBP) levels in germ-free (GF) animals **(b)** Heatmap comparing IEC (CD45-, EpCAM+) gene expression in conventionally raised (CONV-R) vs GF mice. Expression data is relativized to CONV-R-Con mice. # p<0.05 main effect of GF status.

**Figure S5. (a)** Gene expression of toll like receptors (TLR) −4, and −5 do not differ in IECs from germ-free mice as a result of Str or *ex vivo* bacterial (LPS or FLG) challenge. **(b)** Heatmap representing relative gene expression of IECs from CONV-R mice exposed *in vivo* stress (Str) and *ex vivo* bacterial (LPS/FLG) challenge (n=5/group). See Table S1 (GF) and Table S2 (CONV-R) for all means ± SEM and p-values for *ex vivo* experiments.

**Table S1.** Relative expression of select genes within IECs isolated from germ-free (GF) mice exposed to Str paradigm and stimulated ex vivo with LPS or FLG. *p<0.05, **p<0.01.

| Gene | No Stress<br>Mean Relative Expression<br>(+/- SEM) |  |  | Stress<br>Mean Relative Expression<br>(+/- SEM) |  |  | P-value |  |  |  |  |
| --- | --- | --- | --- | --- | --- | --- | --- | --- | --- | --- | --- |
|  | Baseline | LPS | Flagellin | Baseline | LPS | Flagellin | Str<br>Main<br>Effect | LPS<br>Main<br>Effect | FLG<br>Main<br>Effect | Str x<br>LPS | Str x<br>FLG |
| *Duox2* | 1.03<br>(0.04) | 2.21<br>(0.10) | 2.18<br>(0.06) | 1.14<br>(0.03) | 2.76<br>(0.13) | 3.96<br>(0.11) | <0.01 | <0.01 | <0.01 | 0.27 | <0.01** |
| *Nos2* | 1.10<br>(0.18) | 1.65<br>(0.18) | 1.47<br>(0.20) | 1.20<br>(0.27) | 2.75<br>(0.18) | 2.10<br>(0.29) | <0.05 | <0.01 | 0.10 | <0.01** | 0.71 |
| *Fut2* | 0.98<br>(0.06) | 1.15<br>(0.06) | 1.16<br>(0.03) | 0.83<br>(0.09) | 1.30<br>(0.06) | 1.10<br>(0.11) | 0.88 | 0.07 | 0.24 | 0.37 | 0.81 |
| *Saa1* | 1.01<br>(0.03) | 1.68<br>(0.09) | 1.36<br>(0.06) | 0.74<br>(0.04) | 0.92<br>(0.09) | 0.99<br>(0.05) | 0.12 | 0.01 | <0.01 | 0.26 | 0.99 |
| *Il18* | 1.02<br>(0.04) | 0.95<br>(0.03) | 0.73<br>(0.02) | 0.86<br>(0.07) | 0.70<br>(0.13) | 0.68<br>(0.21) | <0.05 | 0.16 | 0.08 | 0.39 | 0.66 |
| *Reg3b* | 1.17<br>(0.09) | 1.14<br>(0.03) | 1.00<br>(0.03) | 0.85<br>(0.06) | 1.15<br>(0.06) | 1.27<br>(0.09) | 0.94 | 0.41 | 0.51 | 0.31 | 0.13 |

**Table S2.** Relative expression of select genes within IECs isolated from conventionally raised (CONV-R) mice exposed to Str paradigm and stimulated ex vivo with LPS or FLG. *p<0.05, **p<0.01.

| Gene | No Stress<br>Mean Relative Expression<br>(+/- SEM) |  |  | Stress<br>Mean Relative Expression<br>(+/- SEM) |  |  | P-value |  |  |  |  |
| --- | --- | --- | --- | --- | --- | --- | --- | --- | --- | --- | --- |
|  | Baseline | LPS | Flagellin | Baseline | LPS | Flagellin | Str<br>Main<br>Effect | LPS<br>Main<br>Effect | FLG<br>Main<br>Effect | Str x<br>LPS | Str x<br>FLG |
| *Duox2* | 1.09<br>(0.16) | 1.38<br>(0.15) | 1.44<br>(0.25) | 1.80<br>(0.18) | 2.03<br>(0.35) | 2.30<br>(0.28) | <0.01** | 0.20 | <0.05* | 0.85 | 0.39 |
| *Nos2* | 1.27<br>(0.18) | 2.30<br>(0.31) | 2.24<br>(0.29) | 2.37<br>(0.61) | 4.07<br>(0.46) | 3.64<br>(0.77) | <0.01** | <0.01** | 0.06 | 0.71 | 0.66 |
| *Fut2* | 1.02<br>(0.09) | 1.01<br>(0.05) | 1.16<br>(0.09) | 1.38<br>(0.17) | 1.76<br>(0.40) | 1.28<br>(0.34) | <0.01** | 0.37 | 0.70 | 0.41 | 0.72 |
| *Saa1* | 1.28<br>(0.29) | 1.09<br>(0.27) | 1.28<br>(0.19) | 1.58<br>(0.30) | 1.89<br>(0.46) | 1.82<br>(0.25) | <0.05* | 0.79 | 0.80 | 0.55 | 0.76 |
| *Il18* | 1.02<br>(0.05) | 1.00<br>(0.14) | 0.92<br>(0.10) | 0.87<br>(0.03) | 1.02<br>(0.09) | 0.82<br>(0.05) | 0.28 | 0.30 | 0.60 | 0.56 | 0.73 |
| *Reg3b* | 1.22<br>(0.57) | 1.81<br>(0.82) | 3.17<br>(1.09) | 3.50<br>(1.18) | 3.84<br>(0.70) | 2.41<br>(0.70) | <0.05* | 0.59 | 0.52 | 0.86 | 0.47 |

**Table S3.**
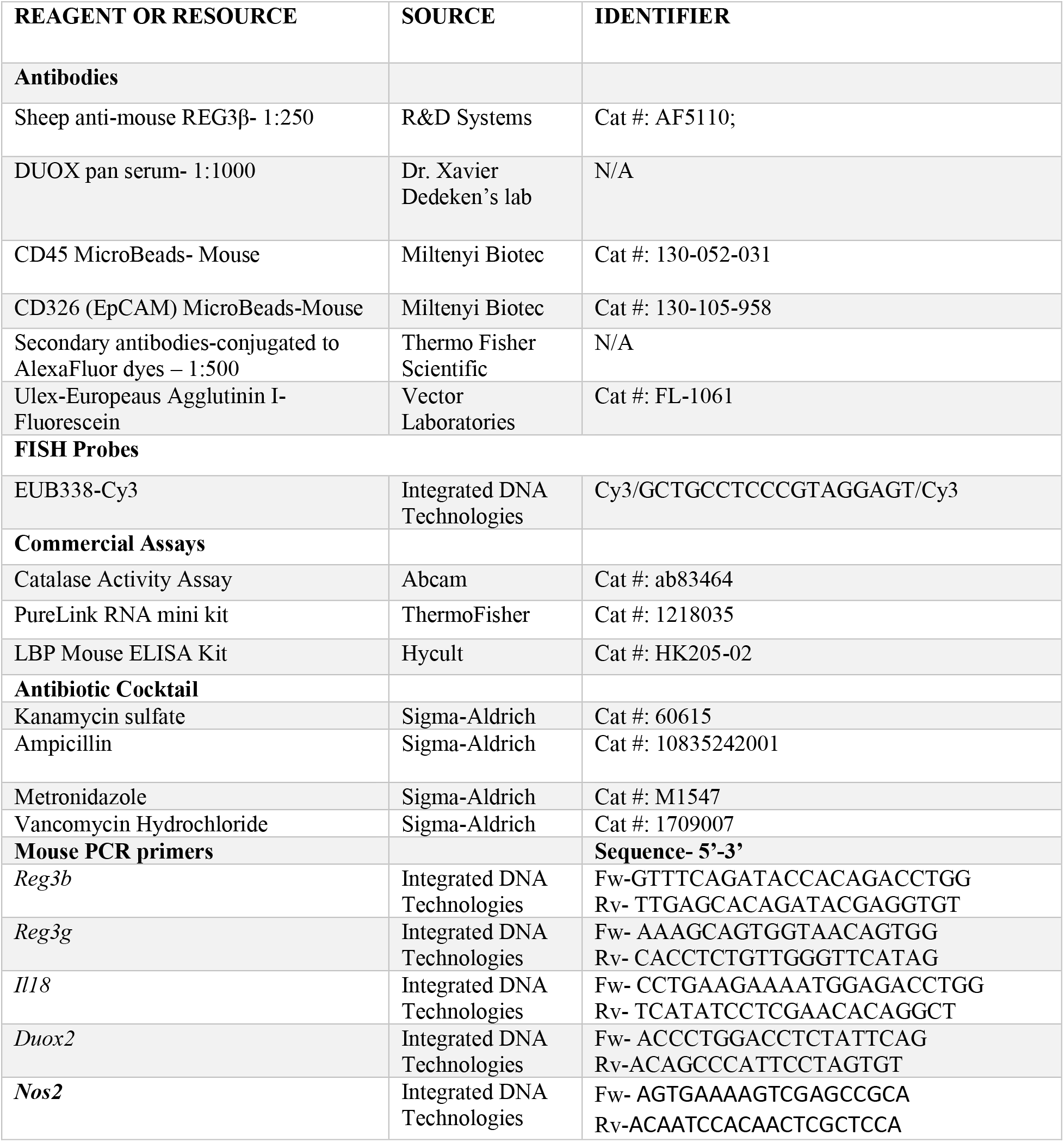
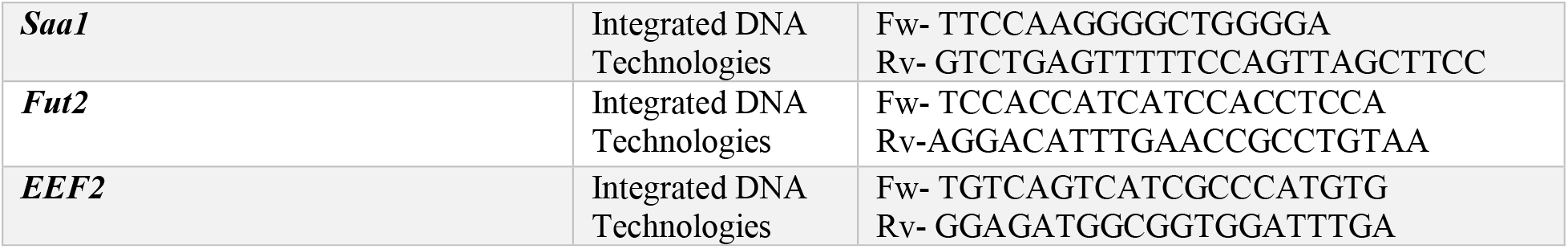
List of reagents and chemicals.

## References

1. Mawdsley JE, Rampton DS. Psychological stress in IBD: new insights into pathogenic and therapeutic implications. Gut. 2005;54(10):1481–91.

2. Goodhand JR, Wahed M, Mawdsley JE, Farmer AD, Aziz Q, Rampton DS. Mood disorders in inflammatory bowel disease: relation to diagnosis, disease activity, perceived stress, and other factors. Inflamm Bowel Dis. 2012;18(12):2301–9.

3. Bailey MT, Dowd SE, Parry NM, Galley JD, Schauer DB, Lyte M. Stressor exposure disrupts commensal microbial populations in the intestines and leads to increased colonization by Citrobacter rodentium. Infect Immun. 2010;78(4):1509–19.

4. Peter J, Fournier C, Durdevic M, Knoblich L, Keip B, Dejaco C, et al. A Microbial Signature of Psychological Distress in Irritable Bowel Syndrome. Psychosom Med. 2018;80(8):698–709.

5. Thaiss CA, Zmora N, Levy M, Elinav E. The microbiome and innate immunity. Nature. 2016;535(7610):65–74.

6. Adolph TE, Tomczak MF, Niederreiter L, Ko HJ, Bock J, Martinez-Naves E, et al. Paneth cells as a site of origin for intestinal inflammation. Nature. 2013;503(7475):272–6.

7. Pickard JM, Maurice CF, Kinnebrew MA, Abt MC, Schenten D, Golovkina TV, et al. Rapid fucosylation of intestinal epithelium sustains host-commensal symbiosis in sickness. Nature. 2014;514(7524):638–41.

8. Schiering C, Wincent E, Metidji A, Iseppon A, Li Y, Potocnik AJ, et al. Feedback control of AHR signalling regulates intestinal immunity. Nature. 2017;542(7640):242–5.

9. Mackner LM, Hatzakis E, Allen JM, Davies RH, Kim SC, Maltz RM, et al. Fecal microbiota and metabolites are distinct in a pilot study of pediatric Crohn’s disease patients with higher levels of perceived stress. Psychoneuroendocrinology. 2020;111:104469.

10. Bailey MT, Dowd SE, Galley JD, Hufnagle AR, Allen RG, Lyte M. Exposure to a social stressor alters the structure of the intestinal microbiota: implications for stressor-induced immunomodulation. Brain Behav Immun. 2011;25(3):397–407.

11. Bailey MT, Lubach GR, Coe CL. Prenatal stress alters bacterial colonization of the gut in infant monkeys. J Pediatr Gastroenterol Nutr. 2004;38(4):414–21.

12. Arora P, Moll JM, Andersen D, Workman CT, Williams AR, Kristiansen K, et al. Body fluid from the parasitic worm Ascaris suum inhibits broad-acting pro-inflammatory programs in dendritic cells. Immunology. 2020;159(3):322–34.

13. Nowarski R, Jackson R, Gagliani N, de Zoete MR, Palm NW, Bailis W, et al. Epithelial IL-18 Equilibrium Controls Barrier Function in Colitis. Cell. 2015;163(6):1444–56.

14. Pircalabioru G, Aviello G, Kubica M, Zhdanov A, Paclet MH, Brennan L, et al. Defensive Mutualism Rescues NADPH Oxidase Inactivation in Gut Infection. Cell Host Microbe. 2016;19(5):651–63.

15. Grasberger H, Gao J, Nagao-Kitamoto H, Kitamoto S, Zhang M, Kamada N, et al. Increased Expression of DUOX2 Is an Epithelial Response to Mucosal Dysbiosis Required for Immune Homeostasis in Mouse Intestine. Gastroenterology. 2015;149(7):1849–59.

16. Huang B, Chen Z, Geng L, Wang J, Liang H, Cao Y, et al. Mucosal Profiling of Pediatric-Onset Colitis and IBD Reveals Common Pathogenics and Therapeutic Pathways. Cell. 2019;179(5):1160–76 e24.

17. Keestra-Gounder AM, Byndloss MX, Seyffert N, Young BM, Chavez-Arroyo A, Tsai AY, et al. NOD1 and NOD2 signalling links ER stress with inflammation. Nature. 2016;532(7599):394–7.

18. Haberman Y, Tickle TL, Dexheimer PJ, Kim MO, Tang D, Karns R, et al. Pediatric Crohn disease patients exhibit specific ileal transcriptome and microbiome signature. J Clin Invest. 2014;124(8):3617–33.

19. Mackos AR, Allen JM, Kim E, Ladaika CA, Gharaibeh RZ, Moore C, et al. Mice Deficient in Epithelial or Myeloid Cell Ikappakappabeta Have Distinct Colonic Microbiomes and Increased Resistance to Citrobacter rodentium Infection. Front Immunol. 2019;10:2062.

20. Mackos AR, Galley JD, Eubank TD, Easterling RS, Parry NM, Fox JG, et al. Social stress-enhanced severity of Citrobacter rodentium-induced colitis is CCL2-dependent and attenuated by probiotic Lactobacillus reuteri. Mucosal Immunol. 2016;9(2):515–26.

21. Galley JD, Parry NM, Ahmer BMM, Fox JG, Bailey MT. The commensal microbiota exacerbate infectious colitis in stressor-exposed mice. Brain Behav Immun. 2017;60:44–50.

22. Zhou G, Soufan O, Ewald J, Hancock REW, Basu N, Xia J. NetworkAnalyst 3.0: a visual analytics platform for comprehensive gene expression profiling and meta-analysis. Nucleic Acids Res. 2019;47(W1):W234–W41.

23. Johansson ME. Mucus layers in inflammatory bowel disease. Inflamm Bowel Dis. 2014;20(11):2124–31.

24. Johansson ME, Phillipson M, Petersson J, Velcich A, Holm L, Hansson GC. The inner of the two Muc2 mucin-dependent mucus layers in colon is devoid of bacteria. Proc Natl Acad Sci U S A. 2008;105(39):15064–9.

25. Lipinski S, Till A, Sina C, Arlt A, Grasberger H, Schreiber S, et al. DUOX2-derived reactive oxygen species are effectors of NOD2-mediated antibacterial responses. J Cell Sci. 2009;122(Pt 19):3522–30.

26. Allen JM, Jaggers RM, Solden LM, Loman BR, Davies RH, Mackos AR, et al. Dietary Oligosaccharides Attenuate Stress-Induced Disruptions in Immune Reactivity and Microbial B-Vitamin Metabolism. Front Immunol. 2019;10:1774.

27. Gautam A, Kumar R, Chakraborty N, Muhie S, Hoke A, Hammamieh R, et al. Altered fecal microbiota composition in all male aggressor-exposed rodent model simulating features of post-traumatic stress disorder. J Neurosci Res. 2018;96(7):1311–23.

28. Bharwani A, Mian MF, Foster JA, Surette MG, Bienenstock J, Forsythe P. Structural & functional consequences of chronic psychosocial stress on the microbiome & host. Psychoneuroendocrinology. 2016;63:217–27.

29. Oshitani N, Kitano A, Okabe H, Nakamura S, Matsumoto T, Kobayashi K. Location of superoxide anion generation in human colonic mucosa obtained by biopsy. Gut. 1993;34(7):936–8.

30. Wilkins TD, Wagner DL, Veltri BJ, Jr., Gregory EM. Factors affecting production of catalase by Bacteroides. J Clin Microbiol. 1978;8(5):553–7.

31. Christian LM, Galley JD, Hade EM, Schoppe-Sullivan S, Kamp Dush C, Bailey MT. Gut microbiome composition is associated with temperament during early childhood. Brain Behav Immun. 2015;45:118–27.

32. Sun Y, Li L, Xie R, Wang B, Jiang K, Cao H. Stress Triggers Flare of Inflammatory Bowel Disease in Children and Adults. Front Pediatr. 2019;7:432.

33. Johansson ME, Sjovall H, Hansson GC. The gastrointestinal mucus system in health and disease. Nat Rev Gastroenterol Hepatol. 2013;10(6):352–61.

34. Kulkarni DH, Gustafsson JK, Knoop KA, McDonald KG, Bidani SS, Davis JE, et al. Goblet cell associated antigen passages support the induction and maintenance of oral tolerance. Mucosal Immunol. 2020;13(2):271–82.

35. Atarashi K, Tanoue T, Ando M, Kamada N, Nagano Y, Narushima S, et al. Th17 Cell Induction by Adhesion of Microbes to Intestinal Epithelial Cells. Cell. 2015;163(2):367–80.

36. Wang Y, Yin Y, Chen X, Zhao Y, Wu Y, Li Y, et al. Induction of Intestinal Th17 Cells by Flagellins From Segmented Filamentous Bacteria. Front Immunol. 2019;10:2750.

37. Andresen L, Jorgensen VL, Perner A, Hansen A, Eugen-Olsen J, Rask-Madsen J. Activation of nuclear factor kappaB in colonic mucosa from patients with collagenous and ulcerative colitis. Gut. 2005;54(4):503–9.

38. Ramakrishnan SK, Zhang H, Ma X, Jung I, Schwartz AJ, Triner D, et al. Intestinal non-canonical NFkappaB signaling shapes the local and systemic immune response. Nat Commun. 2019;10(1):660.

39. Wrzosek L, Miquel S, Noordine ML, Bouet S, Joncquel Chevalier-Curt M, Robert V, et al. Bacteroides thetaiotaomicron and Faecalibacterium prausnitzii influence the production of mucus glycans and the development of goblet cells in the colonic epithelium of a gnotobiotic model rodent. BMC Biol. 2013;11:61.

40. Png CW, Linden SK, Gilshenan KS, Zoetendal EG, McSweeney CS, Sly LI, et al. Mucolytic bacteria with increased prevalence in IBD mucosa augment in vitro utilization of mucin by other bacteria. Am J Gastroenterol. 2010;105(11):2420–8.

41. Parikh K, Antanaviciute A, Fawkner-Corbett D, Jagielowicz M, Aulicino A, Lagerholm C, et al. Colonic epithelial cell diversity in health and inflammatory bowel disease. Nature. 2019;567(7746):49–55.

42. Pereira FC, Wasmund K, Cobankovic I, Jehmlich N, Herbold CW, Lee KS, et al. Rational design of a microbial consortium of mucosal sugar utilizers reduces Clostridiodes difficile colonization. Nat Commun. 2020;11(1):5104.

43. Lafuse WP, Gearinger R, Fisher S, Nealer C, Mackos AR, Bailey MT. Exposure to a Social Stressor Induces Translocation of Commensal Lactobacilli to the Spleen and Priming of the Innate Immune System. J Immunol. 2017;198(6):2383–93.

44. Rendon JL, Li X, Akhtar S, Choudhry MA. Interleukin-22 modulates gut epithelial and immune barrier functions following acute alcohol exposure and burn injury. Shock. 2013;39(1):11–8.

45. Ratsimandresy RA, Indramohan M, Dorfleutner A, Stehlik C. The AIM2 inflammasome is a central regulator of intestinal homeostasis through the IL-18/IL-22/STAT3 pathway. Cell Mol Immunol. 2017;14(1):127–42.

46. Hammer AM, Morris NL, Cannon AR, Khan OM, Gagnon RC, Movtchan NV, et al. Interleukin-22 Prevents Microbial Dysbiosis and Promotes Intestinal Barrier Regeneration Following Acute Injury. Shock. 2017;48(6):657–65.

47. Ivanov, II, Atarashi K, Manel N, Brodie EL, Shima T, Karaoz U, et al. Induction of intestinal Th17 cells by segmented filamentous bacteria. Cell. 2009;139(3):485–98.

48. Ibiza S, Garcia-Cassani B, Ribeiro H, Carvalho T, Almeida L, Marques R, et al. Glial-cell-derived neuroregulators control type 3 innate lymphoid cells and gut defence. Nature. 2016;535(7612):440–3.

49. Grasberger H, El-Zaatari M, Dang DT, Merchant JL. Dual oxidases control release of hydrogen peroxide by the gastric epithelium to prevent Helicobacter felis infection and inflammation in mice. Gastroenterology. 2013;145(5):1045–54.

50. Sommer F, Backhed F. The gut microbiota engages different signaling pathways to induce Duox2 expression in the ileum and colon epithelium. Mucosal Immunol. 2015;8(2):372–9.

51. McCafferty DM. Peroxynitrite and inflammatory bowel disease. Gut. 2000;46(3):436–9.

52. Cross RK, Wilson KT. Nitric oxide in inflammatory bowel disease. Inflamm Bowel Dis. 2003;9(3):179–89.

53. Reinders CI, Herulf M, Ljung T, Hollenberg J, Weitzberg E, Lundberg JO, et al. Rectal mucosal nitric oxide in differentiation of inflammatory bowel disease and irritable bowel syndrome. Clin Gastroenterol Hepatol. 2005;3(8):777–83.

54. Fang FC. Perspectives series: host/pathogen interactions. Mechanisms of nitric oxide-related antimicrobial activity. J Clin Invest. 1997;99(12):2818–25.

55. Kang KA, Ryu YS, Piao MJ, Shilnikova K, Kang HK, Yi JM, et al. DUOX2-mediated production of reactive oxygen species induces epithelial mesenchymal transition in 5-fluorouracil resistant human colon cancer cells. Redox Biol. 2018;17:224–35.

56. Kim SH, Lee WJ. Role of DUOX in gut inflammation: lessons from Drosophila model of gut-microbiota interactions. Front Cell Infect Microbiol. 2014;3:116.

57. Hayes P, Dhillon S, O’Neill K, Thoeni C, Hui KY, Elkadri A, et al. Defects in NADPH Oxidase Genes NOX1 and DUOX2 in Very Early Onset Inflammatory Bowel Disease. Cell Mol Gastroenterol Hepatol. 2015;1(5):489–502.

58. Dhillon SS, Mastropaolo LA, Murchie R, Griffiths C, Thoni C, Elkadri A, et al. Higher activity of the inducible nitric oxide synthase contributes to very early onset inflammatory bowel disease. Clin Transl Gastroenterol. 2014;5:e46.

59. Yoon MY, Min KB, Lee KM, Yoon Y, Kim Y, Oh YT, et al. A single gene of a commensal microbe affects host susceptibility to enteric infection. Nat Commun. 2016;7:11606.

60. Yardeni T, Tanes CE, Bittinger K, Mattei LM, Schaefer PM, Singh LN, et al. Host mitochondria influence gut microbiome diversity: A role for ROS. Sci Signal. 2019;12(588).

61. Ravindra Kumar S, Imlay JA. How Escherichia coli tolerates profuse hydrogen peroxide formation by a catabolic pathway. J Bacteriol. 2013;195(20):4569–79.

62. Ormerod KL, Wood DL, Lachner N, Gellatly SL, Daly JN, Parsons JD, et al. Genomic characterization of the uncultured Bacteroidales family S24-7 inhabiting the guts of homeothermic animals. Microbiome. 2016;4(1):36.

63. Galley JD, Nelson MC, Yu Z, Dowd SE, Walter J, Kumar PS, et al. Exposure to a social stressor disrupts the community structure of the colonic mucosa-associated microbiota. BMC Microbiol. 2014;14:189.

64. Golubeva AV, Crampton S, Desbonnet L, Edge D, O’Sullivan O, Lomasney KW, et al. Prenatal stress-induced alterations in major physiological systems correlate with gut microbiota composition in adulthood. Psychoneuroendocrinology. 2015;60:58–74.

65. Love MI, Huber W, Anders S. Moderated estimation of fold change and dispersion for RNA-seq data with DESeq2. Genome Biol. 2014;15(12):550.

66. Bokulich NA, Kaehler BD, Rideout JR, Dillon M, Bolyen E, Knight R, et al. Optimizing taxonomic classification of marker-gene amplicon sequences with QIIME 2’s q2-feature-classifier plugin. Microbiome. 2018;6(1):90.

67. Quast C, Pruesse E, Yilmaz P, Gerken J, Schweer T, Yarza P, et al. The SILVA ribosomal RNA gene database project: improved data processing and web-based tools. Nucleic Acids Res. 2013;41(Database issue):D590–6.

68. Vazquez-Baeza Y, Pirrung M, Gonzalez A, Knight R. EMPeror: a tool for visualizing high-throughput microbial community data. Gigascience. 2013;2(1):16.

69. Lozupone C, Hamady M, Knight R. UniFrac--an online tool for comparing microbial community diversity in a phylogenetic context. BMC Bioinformatics. 2006;7:371.

70. Michel MC, Murphy TJ, Motulsky HJ. New Author Guidelines for Displaying Data and Reporting Data Analysis and Statistical Methods in Experimental Biology. J Pharmacol Exp Ther. 2020;372(1):136–47.

