## Supplementary figures and images for "Psychological stress disrupts intestinal epithelial cell function and mucosal integrity through microbe and host-directed processes"

### Supplementary Figure 1

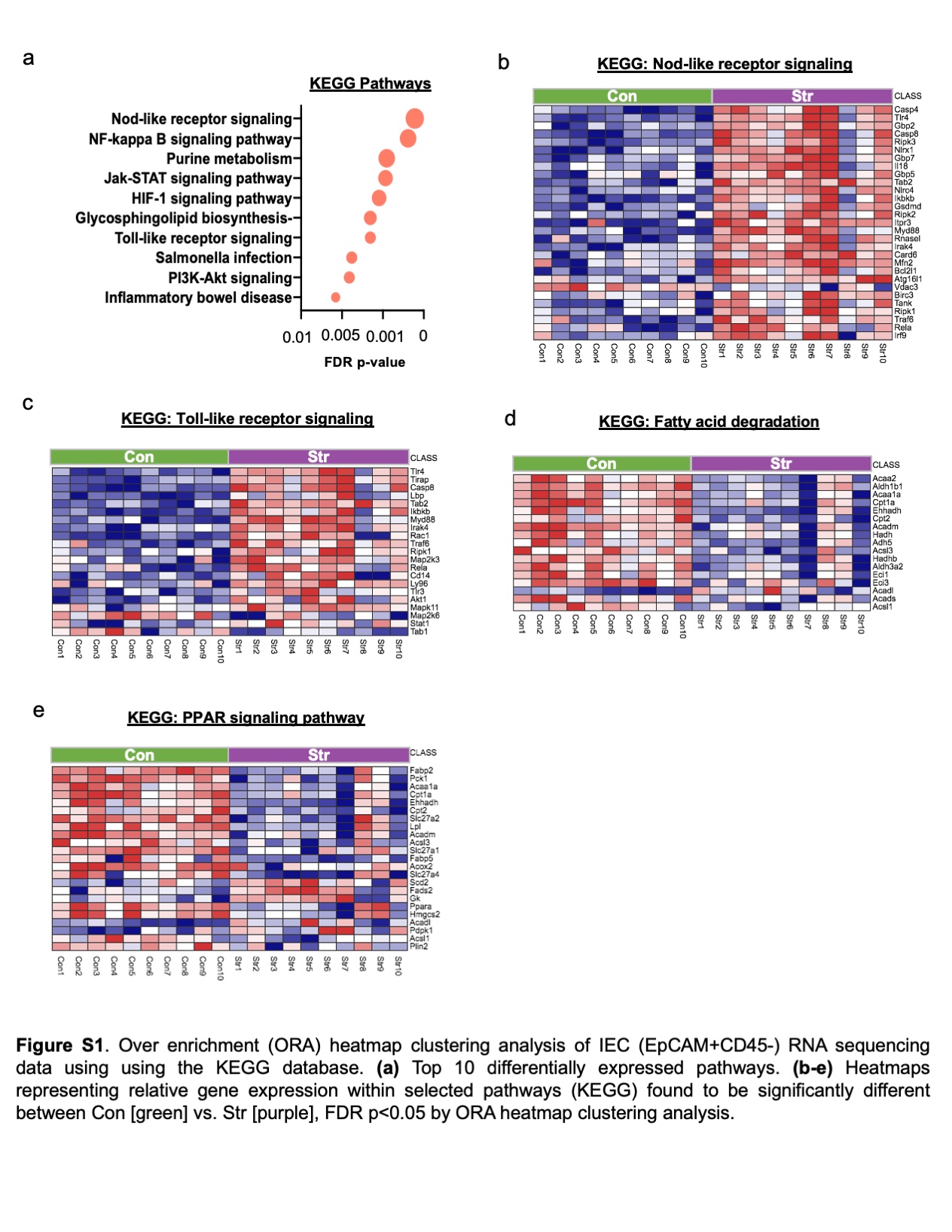

### Supplementary Figure 2

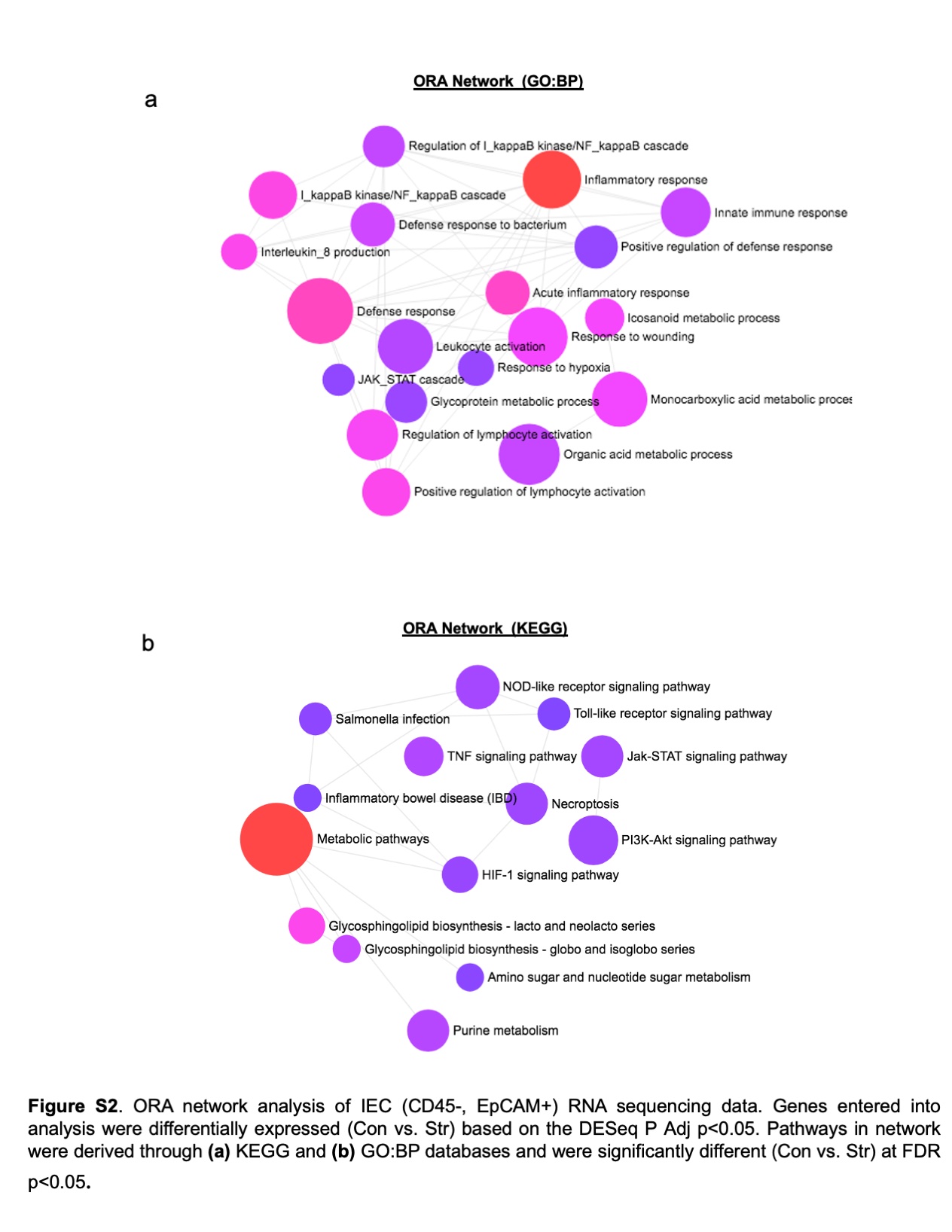

### Supplementary Figure 3

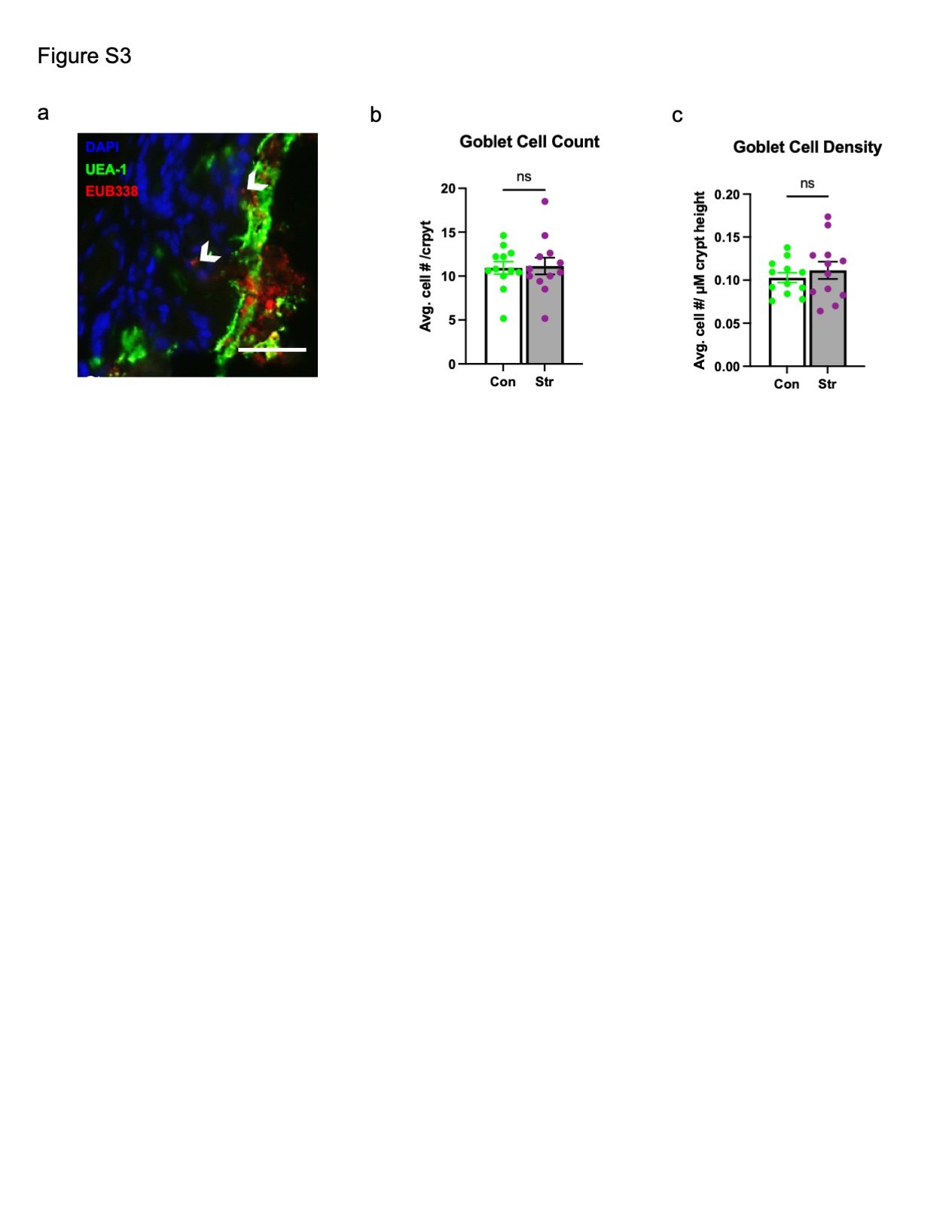

### Supplementary Figure 4

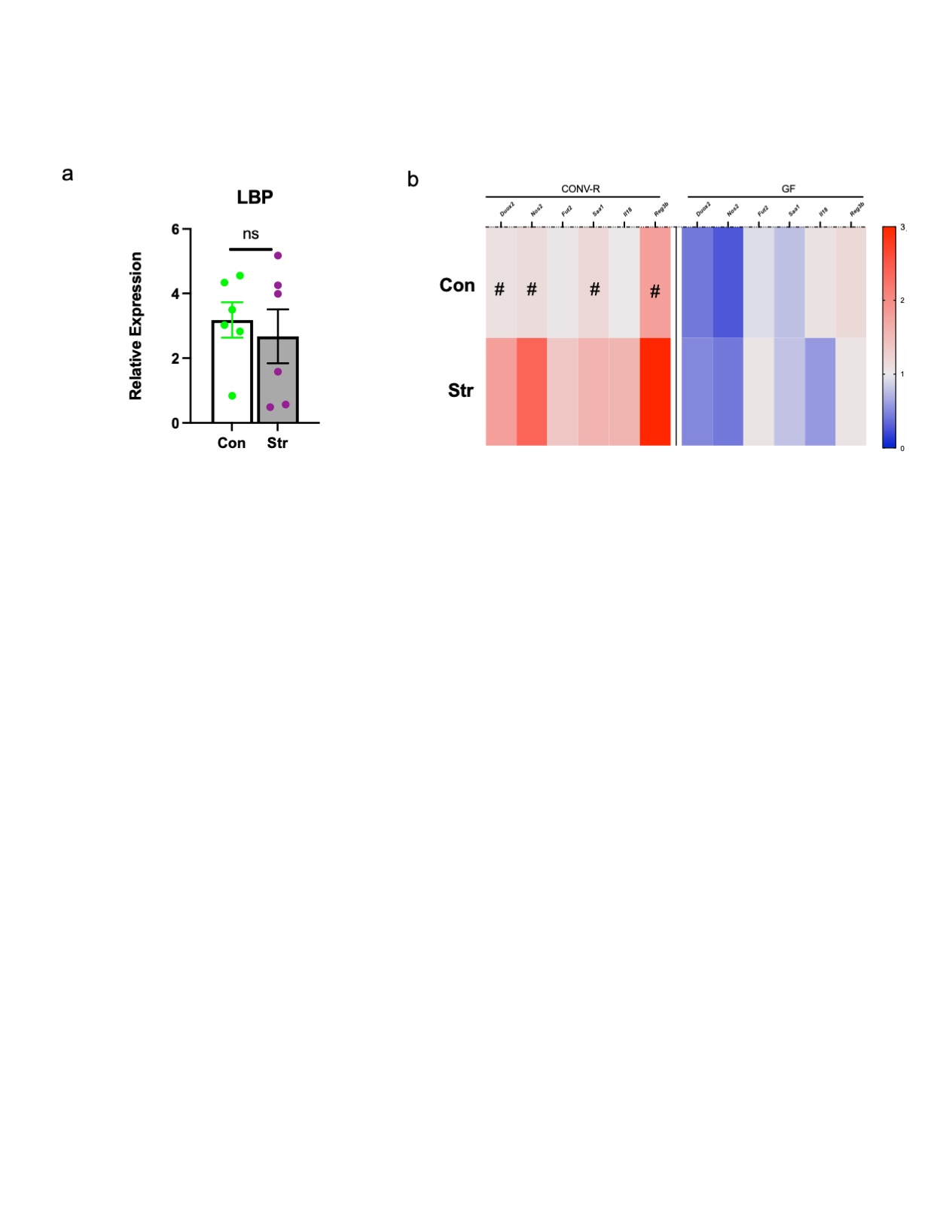

### Supplementary Figure 5

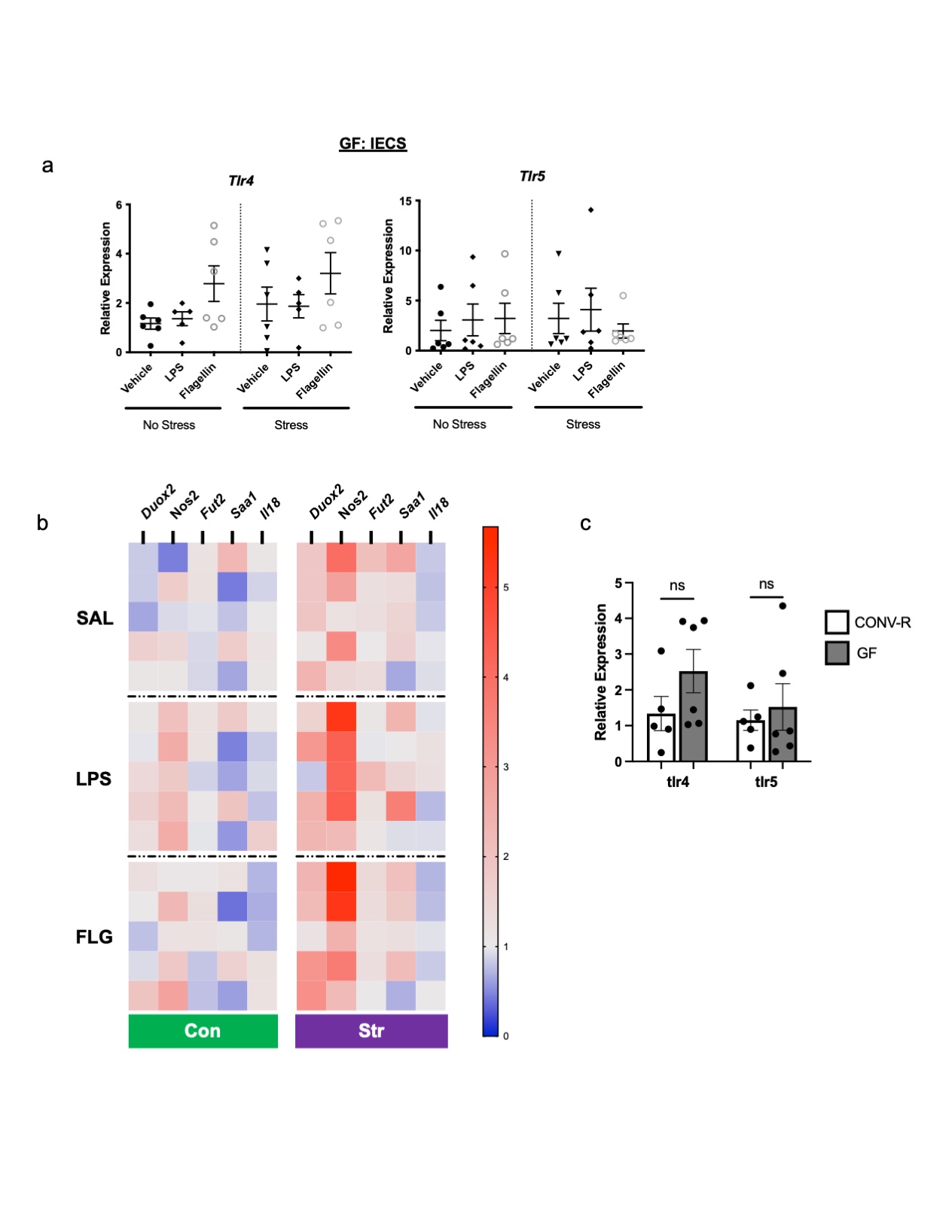
